# MERFISH+, a large-scale, multi-omics spatial technology resolves the molecular holograms of the 3D human developing heart

**DOI:** 10.1101/2025.11.02.686137

**Authors:** Colin Kern, Qingquan Zhang, Yifan Lu, Jacqueline Eschbach, Zehua Zeng, Elie N. Farah, Chu-Yi Tai, Kaifu Yang, Ignatius Jenie, Fenyong Yao, Zoey Zhao, Qixuan Ma, Carlos Garcia Padilla, Alexander Monell, Siavash Moghadami, Fugui Zhu, Bin Li, Angie Hou, Grant Tucker, David Ellison, Neil C. Chi, Xiaojie Qiu, Quan Zhu, Bogdan Bintu

## Abstract

Hybridization-based spatial transcriptomics technologies have advanced our ability to map cellular and subcellular organization in complex tissues. However, existing methods remain constrained in gene coverage, multimodal compatibility, and scalability. Here, we present MERFISH+, an enhanced version of Multiplexed Error-Robust Fluorescence in Situ Hybridization (MERFISH), which integrates chemical probe anchoring in protective hydrogels with high-throughput microfluidics and microscopy. This optimized design supports robust and repeated hybridization cycles across an entire centimeter-scale tissue sample. MERFISH+ allowed to simultaneously quantify over 1,800 genes and resolve the 3D organization of chromatin loci and their associated epigenomic marks in developing human hearts. Using a generative integration framework for spatial multimodal data (Spateo-VI), we harmonized these MERFISH+ transcriptomic and chromatin data to reconstruct a 3D spatially-resolved multi-omic atlas of the developing human heart at subcellular resolution capturing 3.1 million cells across 34 distinct populations. This 3D atlas provides a holistic view of an entire organ enabling the characterization of 3D cellular neighborhoods and transcriptional gradients of substructures such as the descending arteries. Thus, MERFISH+ offers a robust, large-format platform for spatial multi-omics that enables high resolution mapping of gene expression at subcellular resolution and the characterization of cellular organization within 3D organs.

**One Sentence Summary:** MERFISH+ is an spatial multi-omics platform that integrates hydrogel-based probe anchoring, automated high-throughput microfluidics, and large-format multimodal data production to enable comprehensive, subcellular resolution mapping of gene expression and chromatin organization across millions of cells within complex developing human organs.

**Highlights:**

- MERFISH+ expands MERFISH capabilities to measure >1,800 genes and at whole-organ 3D imaging scale
- Combines chemical probe anchoring with high-throughput volumetric microscopy and microfluidics
- Generates a 3D molecular atlas of a developing human heart with > 3.1 million cells at subcellular resolution
- Introduces Spateo-VI, a novel generative framework integrating 3D multimodal datasets

## Introduction

Single-cell RNA sequencing (scRNA-seq) has significantly advanced our understanding of cellular diversity and gene expression dynamics across diverse biological systems ^1–7^. Despite its transformative impact, scRNA-seq inherently disrupts tissue integrity, eliminating critical spatial context. Spatial transcriptomic methods overcome this limitation by preserving positional information, essential for understanding tissue architecture and cell interactions in their native environments ^8–10^. Hybridization-based spatial transcriptomics techniques, such as Multiplexed Error-Robust Fluorescence In Situ Hybridization (MERFISH), Seq-FISH and others^11^ achieve high sensitivity and spatial resolution by repeatedly imaging fluorescently labeled oligonucleotide probes bound to single RNA molecules^12,13^. Many groups have started employing these methods to unravel the spatial gene expression, and cell type and cellular community compositions across an array of human tissues including human heart, kidney, lung, liver and brain^14–19^. Recently, we performed MERFISH in the developing human heart^15^, revealing the spatial organization of single cells into cellular communities that form distinct cardiac structures. However, the current methods face limitations: First, most prior studies focus only on RNA and on a limited set of genes (typically a few hundred), hence overlooking important genes and processes such as epigenetic regulation of DNA which characterize biological processes and diseases^12,13,20,21^. Secondly, most existing studies have focused on imaging limited two-dimensional (2D) tissue sections encompassing up to hundreds of thousands of cells. Although informative, these approaches provide only partial sampling and fail to capture the three-dimensional (3D) architecture of highly complex tissues and organ^20–25^. Many biological processes and cellular interactions occur in three dimensions, especially during a variety of developmental stages in human organogenesis, such as that of the heart ^26^. Therefore, the choice of sectioning plane can significantly influence the observed gene expression patterns and cell organization, potentially leading to biased or incomplete interpretations.

To overcome these limitations, we developed MERFISH+ by extending the capabilities of the MERFISH technology (**Figure S1**). First we incorporated a chemical modification into the encoding/primary probes, covalently anchoring them to a layer of acrylamide gel covering the tissues. These anchored probes are then read out and imaged over an extended period, spanning up to three or more months. This enhanced stability enabled within 2D sections: 1) a new flexible bar-coding scheme allowing for imaging >1,800 genes with a similar quality as that of the 300-gene scale and 2) stable multi-modal imaging of RNA, DNA, and proteins within the same sample. By further combining the stability of the probes with a new high throughput microscope and microfluidics design we increase 10-fold the imaging area, from 1-2 cm^2^ to ~12 cm^2^ allowing for millions of cells profiled per experiment. Extending our previous Spateo framework^26^, we developed a series of novel computational pipelines to facilitate the 3D MERFISH data analyses. We reconstructed a human 3D multi-modal map of the developing heart, revealing the intricate architecture and cellular composition of cardiac 3D neighbourhoods, as well as ligand-receptor interaction flows in three dimensions. Notably, our Spateo-VI method jointly integrated various spatial transcriptomics and spatial multi-omics data to allow the creation of the first transcriptome-wide, multi-modal model of 3D human developing heart, opening doors for many downstream analysis, including the analyses of gene expression and cell-cell communication during coronary artery development.

## Results

### Acrydite FISH-probes improve stability of MERFISH experiments

To overcome the gradual decay of RNA during imaging and hence a gradual decrease of the on-target signal, we modified the MERFISH probe synthesis protocol ^27^ to incorporate an acrydite group to the 5’ end of the oligonucleotides of all the encoding/primary probes. The molecular structure of the acrydite group is similar to the acrylamide monomers and reacts with activated double bonds of acrylamide and bisacrylamide, resulting in the tethering of acrydite-conjugated oligos into the gel matrix ^28^. Such a modification is expected to decouple the probe stability from the decay of the RNA substrate during imaging, greatly expanding the possible number of hybridization cycles (**Figure S1** and **Figure 1A**).

**Figure 1:**
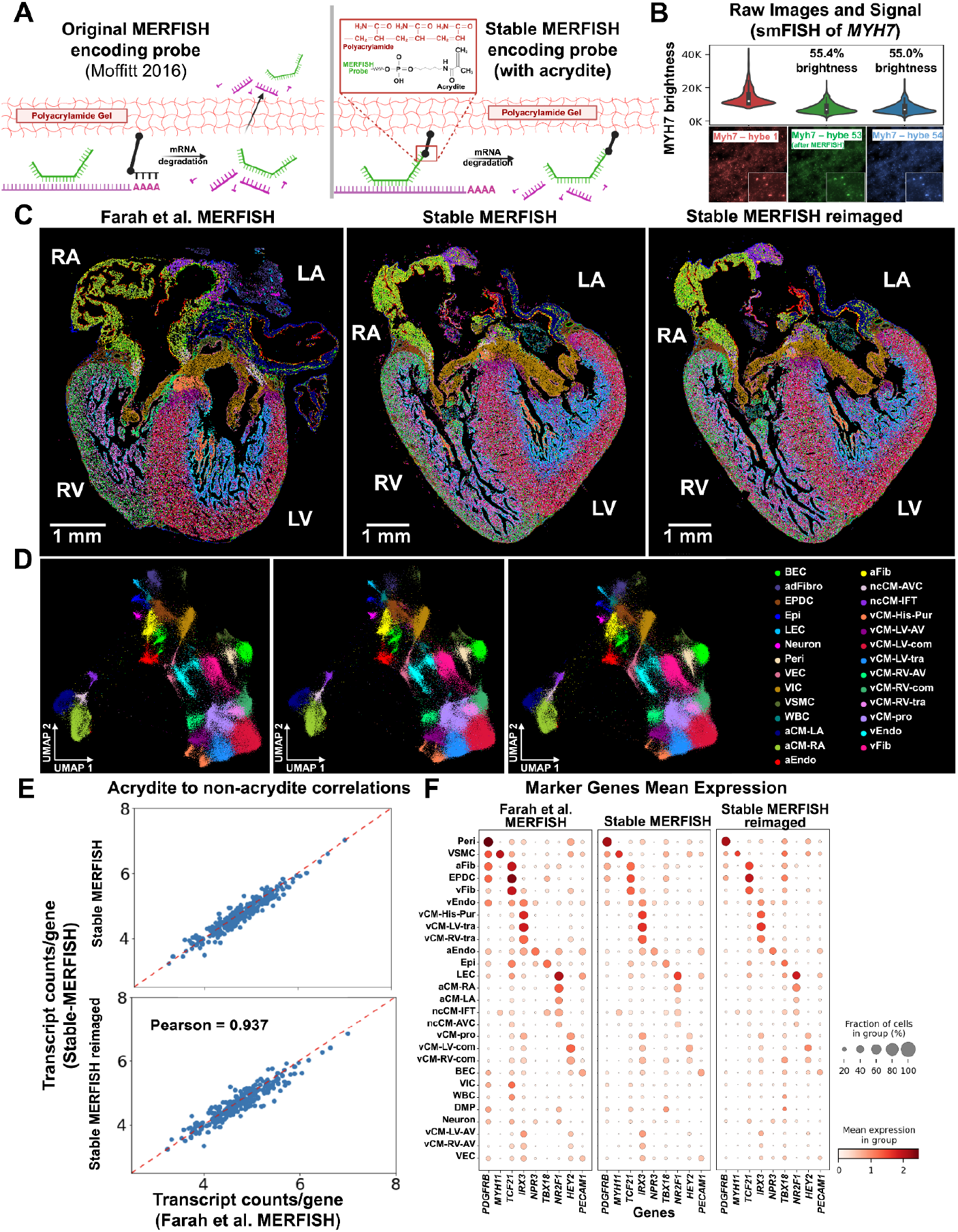
MERFISH+ acrydite modified probes increase stability and robustness of imaging. **(A)** Schematic comparing original and acrydite-conjugated MERFISH probe designs. Left: Original unmodified encoding probes (green) bind to mRNA (purple). A separate acrydite-modified polyT probe hybridizes the polyA tail of mRNA and is linked in the polyacrylamide gel. As mRNA degrades over time, the MERFISH probes become unstable, limiting imaging time. Right: Acrydite-conjugated MERISH+ probes covalently bind directly to the polyacrylamide, enabling probe retention and continued imaging even after mRNA degradation. **(B)** Signal brightness of MERFISH+ probes targeting *MYH7* across 54 cycles of hybridization. Fluorescence intensity plateaus at ~55% of the initial signal observed in round 1. **(C)** Spatial maps of cell types defined across three conditions: original MERFISH probes targeting 238 genes (left), MERFISH+ acrydite-conjugated probes targeting 238 genes (hybridization cycles 1-11 - middle), and reimaged MERFISH+ acrydite-conjugated targeting 238 genes (hybridization cycles 12-22 right). **(D)** UMAP representations of single-cell transcriptional profiles across 238 genes from each experiment for the conditions in (C). Cell types are colored by thei transcriptionally defined cell populations. **(E)** Correlations of average log-transformed gene counts per cell between original and MERFISH+ probes (top, Pearson’s correlation coefficient of 0.959), and between original MERFISH and reimaging MERFISH+ probes (bottom, Pearson’s correlation coefficient of 0.937), showing high concordance. **(F)** Marker gene expression patterns across cell types for the three experimental conditions in (C).

We conducted MERFISH experiments with acrydite modified oligonucleotide probes (called MERFISH+ probes) on 16-um developing human heart sections (12 post-conceptual weeks, 12 pcw). Briefly, a template library of MERFISH probes was synthesized (TWIST Bioscience) and then amplified, first using limited amplification cycles of PCR and then T7 RNA synthesis followed by a reverse transcription reaction with an acrydite modified primer at the 5’ end. These probes were hybridized to RNA molecules within the tissue, followed by the casting of an acrylamide gel onto the sample. In order to quantify the stability of the new probes, we targeted *MYH7* (a gene expressed in ventricular cardiomyocytes) and compared the signal in the first cycle of hybridization with the signal after 54 cycles of hybridization with highly stringent 100%-formamide washes between each cycle. We found that brightness stabilized at about 55% of the first round’s brightness (**Figure 1B**).

We then resynthesized a MERFISH library targeting 238 genes used previously^15^ and performed MERFISH experiments with the acrydite modified probes. On the same sample, we first performed the MERFISH experiment as previously described for 11 hybridization cycles (**Figure 1C, middle**). We then performed a stringent 100%-formamide wash step to completely remove residual readout probes prior to reconducting the MERFISH experiment across an additional 11 hybridization cycles and stringent washes (**Figure 1C, right**). We quantified the number of mRNA molecules of each of the 238 targeted genes in each cell and performed UMAP embedding^15^ and cell type definition using Leiden clustering of the cells imaged across the two MERFISH experiments. We obtained a near-identical cell type definition and spatial distribution across the 2D section (**Figures 1C and 1D**) with the same cell type markers defining each cell type (**Figure 1F**) as in our prior publication with the original MERFISH protocol^20^. Furthermore, we obtained a very high correlation (Pearson’s correlation coefficients of 0.93 and 0.95) with near identical average expression per cell using either MERFISH+ or conventional MERFISH probes^20^, suggesting that the detection efficiency and accuracy was not appreciably changed by adding the acrydite modification (**Figure 1E**). We note that in conventional MERFISH experiments, failure can occur due to multiple reasons such as RNA degradation, equipment malfunctions, or instability of imaging reagents. With MERFISH+ probes, however, a stringent 100% formamide wash allows the experiment to be reset and resumed without compromising the sample. Together, these improvements make MERFISH+ probes a more robust and reliable platform for high-resolution spatial transcriptomics, especially for limited human samples.

### MERFISH+ probes allows for scaling up the number of target genes

The increased stability of the MERFISH+ probes allow for a scalable increase in the number of genes imaged to thousands of targets (**Figures S2 and S3**). We imaged a 12 p.c.w. heart section across a total of 1,835 genes. These genes were selected based on prior snRNAseq to allow for a high definition of cell types across multiple tissues including heart^15^, brain^29^, pancreas^30^ and lung^31^. Notably, 51% (935) of these genes also showed high differential expression in heart scRNA-seq data^15^. We imaged these genes using a total of 112 cycles of hybridization across 7 modules of 16 hybridization cycles each (**Figure S2**). As controls, we imaged across two of the modules all the previously profiled 238 genes twice using two combinatorial encoding strategies obtaining high correlation with previous measurements (**Figures S2G and S2H**). Cell type identities were determined de novo based on clustering of the single-cell expression profiles across the 1,835 genes, and exhibited strong concordance with matching populations identified by scRNA-seq^15^ (**Figures 2A and 2B**). With an increased number of profiled genes, we identified a few additional transcriptionally distinct cell populations compared to our prior work^15^ (**Figure 2A-boxed**). First, we highlight an additional cardiac cell subcluster we named vCM-LV-papillary (**Figures 2C-yellow**) that was anatomically located within the papillary muscle regions of the left ventricle between the ventricular wall and valvular tissues (**Figure 2C**). In addition to expressing inner chamber cardiomyocyte markers (i.e. *IRX3*) (**Figure S4A**), these cells also expressed specifically *DLGAP1* and *ACTA1*, distinguishing them from other vCMs (**Figures 2C-right, 2D and S1**).

**Figure 2:**
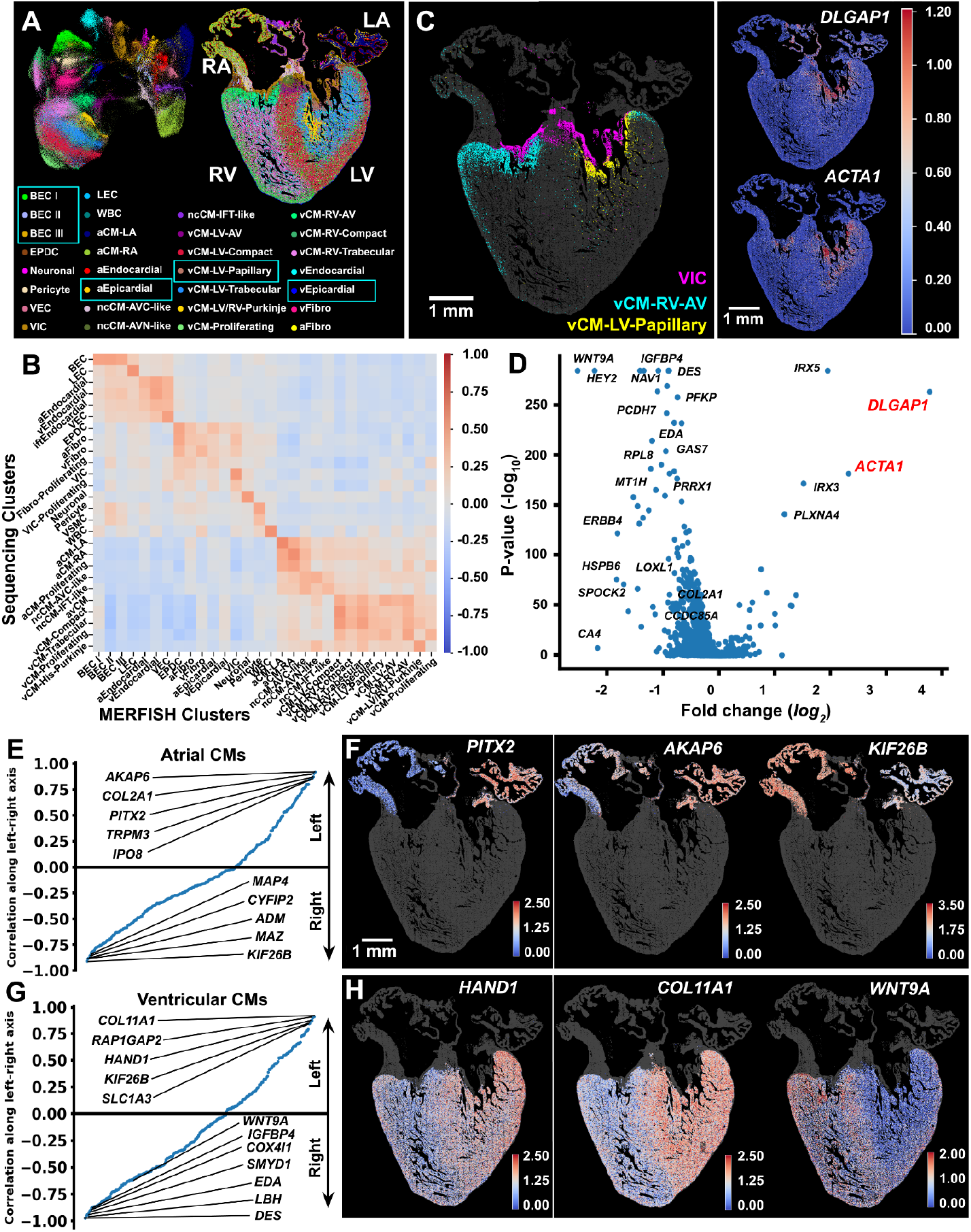
MERFISH+ probes allow for imaging >1,800 genes in human heart tissue. (**A**) UMAP and spatial visualization of cell clusters identified in a 12 p.c.w. human heart using MERFISH+ probes targeting 1,835 genes. Newly identified cell populations (compared to^15^) are highlighted by boxes, including a subcluster of cardiomyocytes (vCM-LV-Papillary), three subclusters of blood endothelial cells (BEC I, II, III), and two subclusters of epicardial cells (aEpicardial and vEpicardial). The spatial distribution of the cell populations identified is shown with matching colors - right. (**B**) Correlation matrix of average gene expression for 1,836 genes between MERFISH+ cell-type clusters and sequencing-derived clusters^15^. (**C**) Left: Spatial distribution of the newly identified vCM-LV-Papillary cluster (yellow), together with the spatial distribution of VIC (magenta) and vCM-RV-AV (teal) clusters for anatomical reference. Right: Spatial gene expression pattern of vCM-LV-Papillary markers *DLGAP1* and *ACTA1*. (**D**) Volcano plot of differential gene expression between vCM-LV-Papillary and other vCM cells. Positive fold change indicates higher expression in vCM-LV-Papillary. X-axis marks log_2_ fold change and Y-axis marks –log_10_ p-value. (**E**) Correlation of gene expression with gradient along the lef-right axis for atrial cardiomyocytes (aCMs). (**F**) Spatial expression pattern for selected genes in aCMs with left-right gradients including *PITX2* (canonical LA marker [reference]), *AKAP6*, and *KIF26B*. (**G**) Correlation of gene expression with gradient along the lef-right axis for ventricular cardiomyocytes (vCMs). (**H**) Spatial expression pattern for selected genes in vCMs with left-right gradients for *HAND1* (canonical LV marker), *COL11A1*, and *WNT9A*.

We also identified transcriptionally distinct populations of the blood endothelial cells (BECs), which we named BEC I, II and III (**Figures 2A-boxed, S4A)**. All these cells express genes characteristic of BECs (i.e. *PECAM1,CDH5*) as well as genes differentially expressed in these BECs subtypes (*HEY2* higher in BEC I, *HEYL* and *MSC* higher BEC II and *SOX17* and *DLL4* higher in BEC III) (**Figure S4A and S4C**). Spatially, BEC I is the most abundant and is broadly distributed along the ventricular wall and intraventricular septum (**Figure S4B**). The BEC II endothelial subcluster is located at the sinus venosus region or the atrioventricular canal junction between the atrium and ventricle, consistent with a sinus venosus–derived endothelial lineage (**Figure S4B)**^32^. In contrast, BEC III cells are sparsely distributed within the ventricular myocardium, forming distinct clusters. These cells show high expression of *SOX17*, a key regulator of coronary artery development ^33^, as well as *DLL4*, a NOTCH pathway gene also essential for coronary artery formation^34,35^ (**Figures S4B-S4D**). Similarly, we were able to distinguish transcriptionally the epicardium into two distinct populations: aEpicardial and vEpicardial (**Figure S4E**). These two populations express marker genes of epicardial cells (i.e. *TCF21*) and are specifically located in the atrial surface (aEpicardial, marked by elevated *F13A1* expression) and in the ventricular surface (vEpicardial, marked by elevated *CEMIP* expression) (**Figures S4D and S4E**).

Across the 1,835 genes profiled we also identified genes with asymmetric expression between the left and right sides of the heart (**Figures 2E–2H**). By focusing on atrial and ventricular cardiomyocytes and ordering genes based on their left-right expression gradients (**Figures 2E and 2G**), we confirmed the left-side bias of *PITX2* ^36^, and *HAND1*^*37*^, which are known markers of the left atrium (**Figures 2E and 2F**) and left ventricle, respectively (**Figures 2G and 2H**). In addition, in atrial cardiomyocytes, we identified genes such as *AKAP6, COL2A1, and TRPM3* as enriched in the left atrium, while *ADM* and *KIF26B* were more highly expressed in the right atrium (**Figures 2E and 2F**). Similarly, in ventricular cardiomyocytes, genes such as *COL11A1, RAP1GAP2*, and *KIF26B* were highly enriched in the left ventricle, whereas *SMYD1, EDA, LBH, WNT9A* and *DES* were enriched in the right ventricle (**Figures 2G and 2H**). Notably, several asymmetrically expressed genes, such as *COL11A1*, are essential for ventricular morphogenesis ^38,39^. Together, these results demonstrate a methodology for scaling up the number of genes profiled, providing a more detailed spatial map of additional cell populations and gene expression gradients in the developing human heart.

### Facile multimodal imaging of RNA, DNA and proteins

We next investigated whether the improved stability of MERFISH+ probes facilitates multi-modal imaging of: RNA transcripts, 3D DNA chromatin structure and nuclear accumulation of epigenetic marks (**Figures 3A-3D**). We selected and imaged a genetic locus, the *FXYD* region which contains 21 genes, including the *FXYD* family of ion channel modulators (**Figure 3B)**, which show variability of expression across the cell types of the developing heart^15^ (**Figures 3E**). We synthesized ~24,000 oligonucleotide probes with acrydite anchor, targeting this entire 510 kb *FXYD* locus via DNA FISH. Using sequential hybridization and imaging, we readout the signal of 51 10-kb segments comprising this locus across all the cells in a 12 p.c.w. human heart section. By fitting and aligning the signal of each segment, the 3D structures of the *FXYD* locus in each chromosome in each cell is reconstructed (**Figures 3D and 3F**) with ~50 nm spatial resolution and 10-kb genomic resolution. On a single-cell level, we noticed that this locus tended to organize within globular domain-like structures (**Figure 3D, right**) reminiscent of globular, TAD-like structures previously reported^40^. Upon averaging the inter-distances between each of the 10-kb segments across ~36,000 cardiomyocytes, we found excellent agreement between the average structure determined by imaging and that determined by an orthogonal sequencing method called HiC in cultured cardiomyocytes^41^ (**Figure S5A**). To relate the chromatin structure with transcription within the same cells we further imaged the mature RNA expression of 238 marker genes and both the mature and nascent RNA expression of the genes in the *FXYD* locus (**Figures 3C and 3D)**. The MERFISH+ anchoring scheme facilitated the imaging of RNA across additional cycles after the DNA measurements. The RNA expression was used to define cell types, obtaining similar quality to the experiments in which only RNA is measured (**Figures S5B**), and to quantify the expression of the *FXYD* genes across cell types (**Figures 3E**). We asked if there are specific structures of this locus that are differentially formed across different cell types. By comparing the average inter-distances of the genomic loci comprising FXYD region, we identified a “loop” (closer distance) between the promoter of the *FXYD5* and a distal element close to the promoter regions of *LSR* and *USF2* specifically in endothelial cells compared to all other cell types (**Figure 3G-green arrow**). Intriguingly this long-range chromatin interaction (70 kb) coincides with the increased expression of the *FXYD5* gene within endothelial cells (**Figure 3E)**. Additionally, we also saw a potential compartment switch in which the first chromatin domain (chr19:35.00Mb-35.14Mb) was on average closer with the last chromatin domain (chr19: 35.23Mb-35.50Mb) in ventricular cardiomyocytes compared to all other cell types (**Figure 3H**). Collectively, these results demonstrate that MERFISH+ facilitates high-accuracy tracing of chromatin structures and quantification of RNA expression in single-cells, identifying distinct chromatin organizational patterns across different cell types.

**Figure 3:**
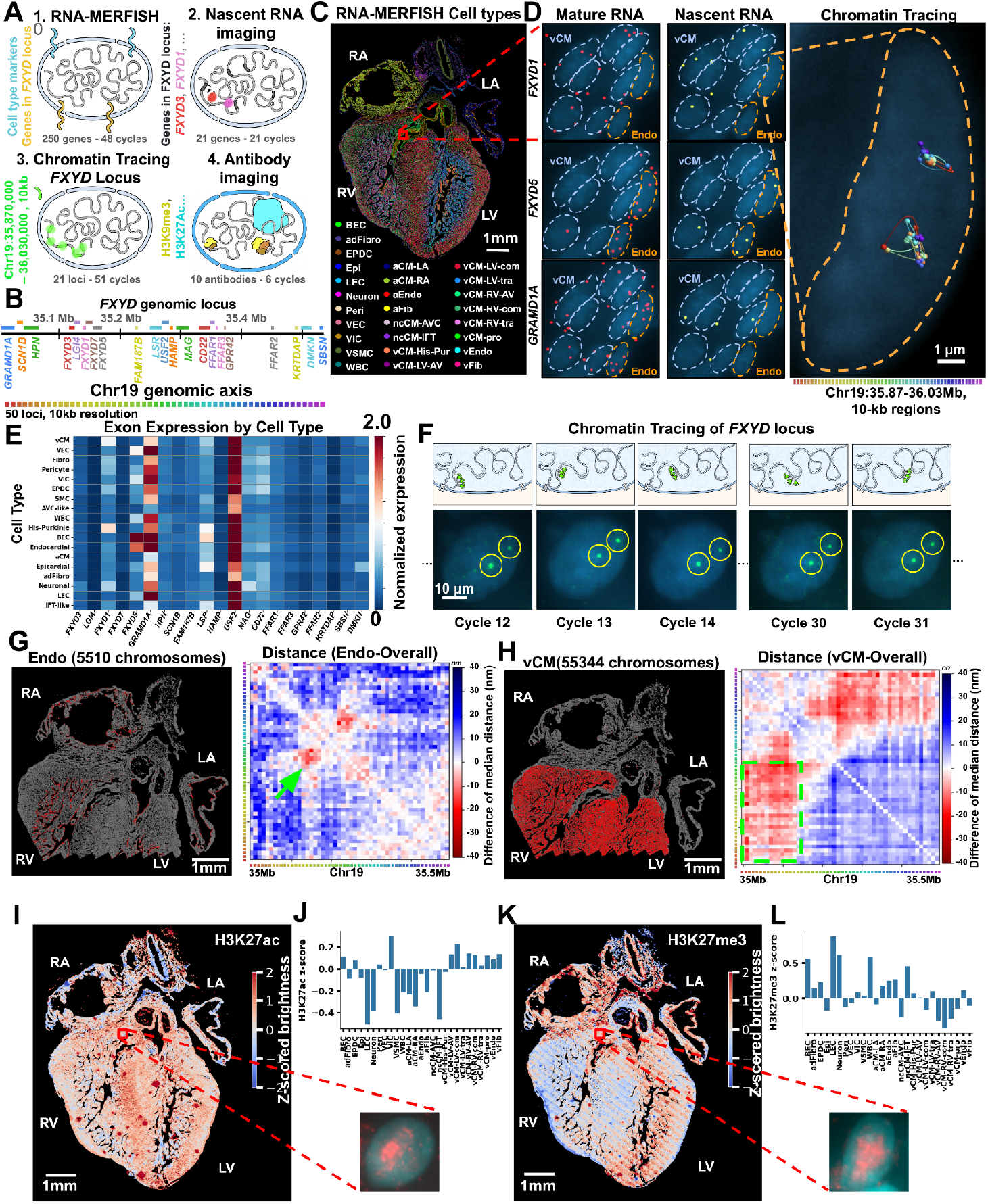
MERFISH+ probes facilitate multimodality imaging of RNA, DNA and epigenetic marks. (**A**) Schematic of multimodal imaging combining RNA MERFISH of mature RNA, nascent RNA, chromatin tracing and epigenetic protein marks. (**B**) FXYD locus (chr19:35Mb-35.51Mb) and comprising genes selected for chromatin tracing imaging at 10 kb genomic resolution. (**C**). MERFISH image (left) of cell type distribution with 27 cell clusters defined based on mature RNA imaging of 238 genes. (**D**) Zoom-in images of mature RNA of genes in the FXYD locus (left), their corresponding nascent RNA (middle) and reconstruction of 3D chromatin structure for two FXYD loci located in chromosome 19 homologues in a single nucleus (right). (**E**) Gene expression matrix across cell types of mature RNA for the genes in the FXYD locus. (**F**) Top: scheme for chromatin tracing of 51 10-kb regions comprising the FXYD locus imaged across 51 cycles of hybridization. Bottom: Example images of the DNA FISH signal across different cycles of hybridization. (**G**) (**H**) Difference of median distance matrices for all the 51 10kb regions of the FXYD locus between endothelial cells and all other cells (G) and between ventricular cardiomyocytes (vCMs) and all other cells (H). Green arrow points to lower distance in endothelial cells between the FXYD5 promoter and a distal element close to the LSR and USF2 promoter. Green box highlights smaller distances in vCMs between the two non-neighboring chromatin domains. (**I**) Spatial map of the z-scored antibody brightness of the H3K27ac mark across cells in the heart section from (B). **(J)** Bar plot of z-scored antibody brightness of the H3K27ac mark across cell types called in (B). (**K**),(**L**) Same as (I), (J) for H3K27me3 mark.

To further test the multimodal capability of the MERFISH imaging and complement our chromatin structure analyses, we performed sequential antibody staining and immunofluorescence imaging on the same sample with different epigenomics markers: pol2ser2, pol2ser5, H3K4me3, H3K27ac, and H3K27me3 (**Figures 3l-L and S5C, S5D**). The total brightness of the antibody signal in each nucleus provided an approximate measure of the accumulation of that mark within each cell allowing for a comparison across cell types. We observed that the patterns of these epigenomic markers exhibited both cell type–specific and spatial differences in a reproducible manner across replicate experiments (**Figures 3I, 3K, S5C and S5D**). H3K27ac, associated with active chromatin regions^42,43^, was correlated with the signal for Pol2Ser2, marking actively elongating RNA polymerase^44^ (**Figure S5E**) and showed higher signal in VIC, vCM-LV-com and vCM-RV-com cells compared to neuronal-like, LEC, ncCM-IFT, aCM-LA, and aCM-RA cell populations. In contrast, H3K27me3, associated with polycomb repressed chromatin^45^, was correlated with H3K9me3 levels (**Figure S5F**), marking mainly constitutive heterochromatin^46^ and was anti-correlated across cell types with the active chromatin mark H3K27ac (**Figure S5G**). Overall, MERFISH+ probes facilitate multi-modal measurements connecting the kb-scale DNA structure, RNA expression and epigenetic states of different cell types.

### High-throughput imaging platform for molecular profiling of an entire organ

To comprehensively capture the cellular composition of the entire heart, broader tissue coverage is required, necessitating spatial methods capable of large-area imaging. We argued that the increased stability of the anchored probes will enable large-scale imaging. We developed a new microscope system developed on top a more flexible Applied Scientific Instrumentation microscope body equipped with higher-power lasers (up to 2.5W), a larger-format camera (20.8 mm sensor), and an upgraded microfluidics chamber, achieving a 10-fold increase in imaging area compared to previous and commercially available MERFISH imaging platforms^15^ (**Figures 4A, 4B and STAR Methods**). We provide a complete list of parts to assemble this high-throughput microfluidics-microscope system. To comprehensively examine the heart, we utilized this system to image 238 genes^15^ across a total of 53 consecutive transverse sections from a 12 p.c.w. human heart donor across two experiments (**Figure 4C**). This imaging scheme covered the entire developing heart at 112 um resolution along the superior-inferior axis, profiling 3.1 million cells with high correlation of average gene expression to previous scRNA-seq and MERFISH data^15^ (**Figures S6A and S6B**).

**Figure 4:**
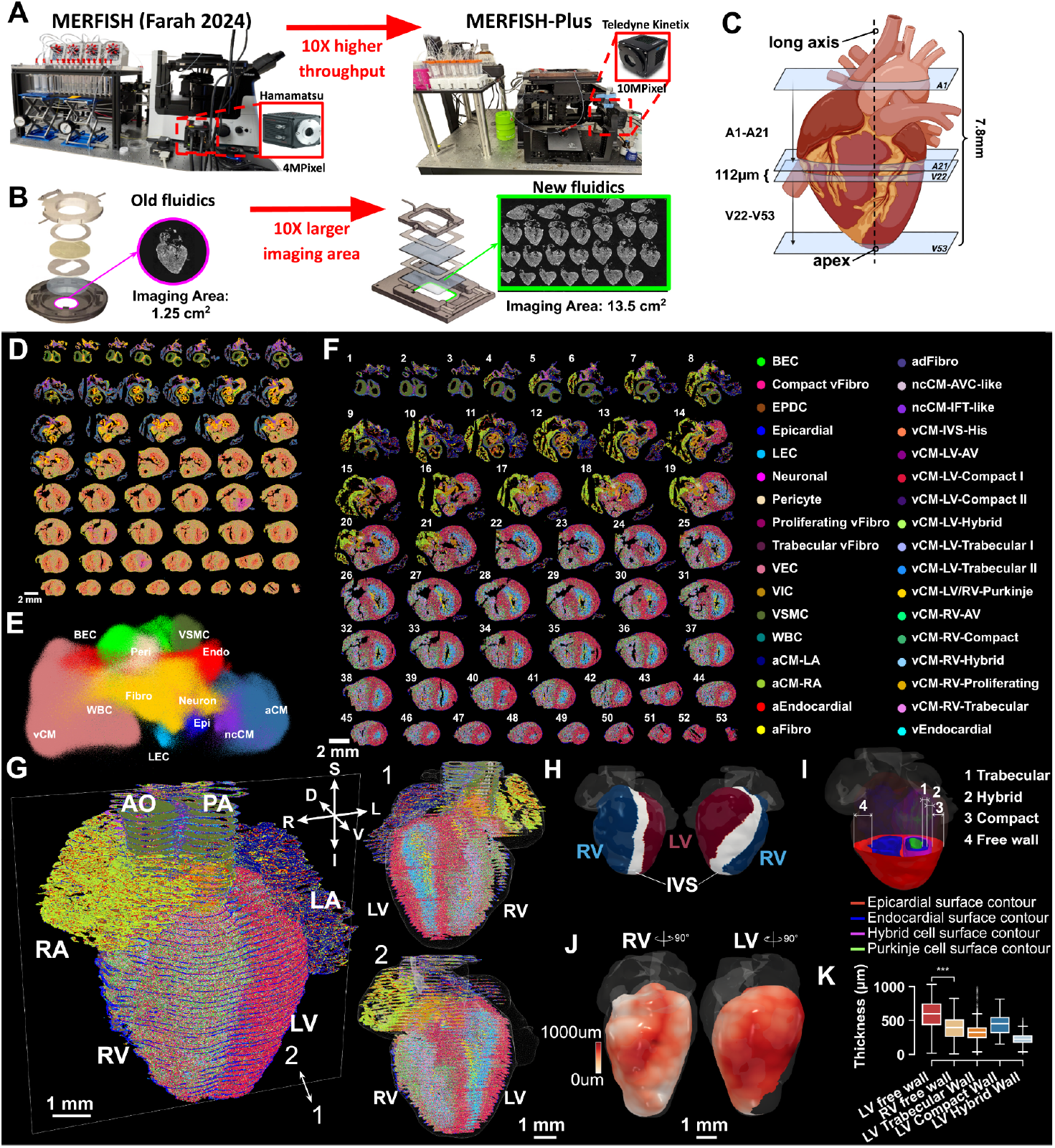
High-throughput MERFISH+ microfluidics/microscopy system enables 3D reconstruction of a molecular hologram of the developing human heart. **(A),(B)** Images of MERFISH+ new microscopy (A) and microfluidics (B) compared to previous MERFISH instrumentation^15^ resulting in 10-fold increase in throughput. (**C**) Schematic of coronal serial sectioning of a human heart donor used for MERFISH+. 53 coronal sections with 112 um inter-spacing were imaged across 2 MERFISH+ experiments. (**D, E**) Spatial distribution of 12 major cell types (D) and corresponding transcriptional UMAP (E) of the 53 sections imaged by MERFISH+. (**F**) Spatial distribution of high resolution cell types with consistent calling as in^15^. **(G)** Left: 3D-reconstruction of an entire heart with 53 sections imaged by MERFISH+ aligned using Spateo^26^. The cell type clusters are colored as in (F). Right: Views of the two halves of 3D-reconstruction bisected along the central left-right plane. These images show the internal spatial distribution of different cell types. **(H)** 3D-images highlighting the left and right ventricular regions used to define the free wall. **(I)** Images of 3D meshes encapsulating the endocardium and epicardium for calculating free wall thickness (4) and 3D meshes encapsulating vCM-LV/RV-Purkinje cells, vCM-LV-Trabecular cells, vCM-LV-Hybrid cells and the vCM-LV-Compact cells for calculating the thickness of the trabecular layer (1), the hybrid layer (2) and the compact layer (3) of the left ventricle in 3D. **(J)** Heatmap of ventricular free wall thickness of the right ventricle (left) and left ventricle (right), across the surface of the heart. Red corresponds to thicker wall and white to thinner wall. **(K)** Box plots of free wall thickness across the left and right ventricles and the trabecular, hybrid and compact layer thicknesses across the left ventricle. Two-sided Student’s t-test was used to compare left vs right free wall thickness. *** indicates a p-value < 1e^−3^.

### Reconstructing a 3D cell atlas of developing human heart

Cell types were defined by label transfer from previous developing heart MERFISH data ^15^, with excellent correspondence for matching cell types (**Figure S6C**). We captured all 12 major cell types in the heart: blood vessel endothelial cells (BEC), atrial cardiomyocytes (aCM), non-chambered cardiomyocytes (ncCM), ventricular cardiomyocytes (vCM), endocardial cells (Endo), epicardial cells (Epi), fibroblasts (Fibro), lymphatic endothelial cells (LEC), pericytes (Peri), white blood cells (WBC), vascular smooth muscle cell (VSMC), and neural-like cells (**Figures 4D and 4E**). At the fine cell type level, the spatial distribution of 34 distinct cell clusters was defined (**Figures 4F, S7**). This comprehensive sampling provides an unbiased assessment of the abundance of the cell populations within the developing human heart (**Figures S6D and S6E**). The number of cells of each cell type varied by two orders of magnitude, ranging from the most abundant cardiomyocyte populations (vCM-LV-com I, 20.45%, with an estimated ~5,000,000 cells within the entire 12 p.c.w. heart) to less abundant cell types such as neuronal-like cells (0.36%, ~89,000 cells) or specific types of endothelial cells (VEC: 0.74%, ~183,000 cells and LEC: 0.47%, 115,000 cells).

Using the recently developed molecular alignment tool, Spateo^26^, we aligned the 53 imaged sections into a 3D reconstruction of the developing human heart at single-cell resolution (**Figure 4G**). This digital model provides the 3D spatial distribution of each cell type (**Figure S7**). In addition, it enables quantitative analysis of 3D anatomical parameters (**STAR Methods**), such as measuring the ventricular free wall thickness defined here as the distance between the epicardial surface contour and the endocardium contour (**Figures 4H-4K)**. Consistent with prior studies^47^, our analysis reveals that the free wall of LV is generally thicker than the RV. Moreover, the wall thickness is not uniform, showing regional variation in both the LV and RV (**Figures 4I-4K, S8A and S8B**) ^47^. Furthermore, the thickness of the different molecular layers of the ventricular wall can be quantified based on the composing cell types. Specifically, for the left ventricle, we defined 3D meshes encapsulating the vCM-LV-Trabecular cell types, vCM-LV-Hybrid cell type and the vCM-LV-Compact cell type and calculated in 3D the regional thickness of the trabecular, hybrid and compact layers (**Figures 4I and 4K**). We found that the hybrid layer was the thinnest and proportionally changed across the heart in a correlated manner with the compact layer thickness (**Figures 4K and S8C**).

### Cellular neighborhoods within the 3D heart

To quantify how cell types spatially associate within the 3D heart, we performed a cellular neighborhood analysis^48^ (**STAR Methods**). Briefly, we quantified for each cell the composition of neighboring cell types and correlated these neighborhood compositions to identify pairs of cell types that are consistently proximal. Clustering these patterns further grouped cell types into five hierarchical spatially-defined cellular neighborhoods **(Figures 5A-5C**). Many of these neighbourhoods were dominated by a few cell types but also captured less abundant cell populations that tend to co-localize with them in 3D. For instance, in neighborhood (I), containing predominantly compact and hybrid ventricular cardiomyocytes, we also found enrichment of blood endothelial cells and pericytes (**Figures 5A and 5B**), indicative of increased blood vessel formation^49^, and a specific subpopulation of fibroblasts (Compact vFibro) consistent with our prior report^15^. Similarly, in the neighborhood (II) enriched in trabecular cell types, we found an enrichment of specific fibroblast and endocardial subpopulations (vFibro and vEndocardial), consistent with the role of endocardium in regulating trabecular cardiomyocyte differentiation^50^.

**Figure 5:**
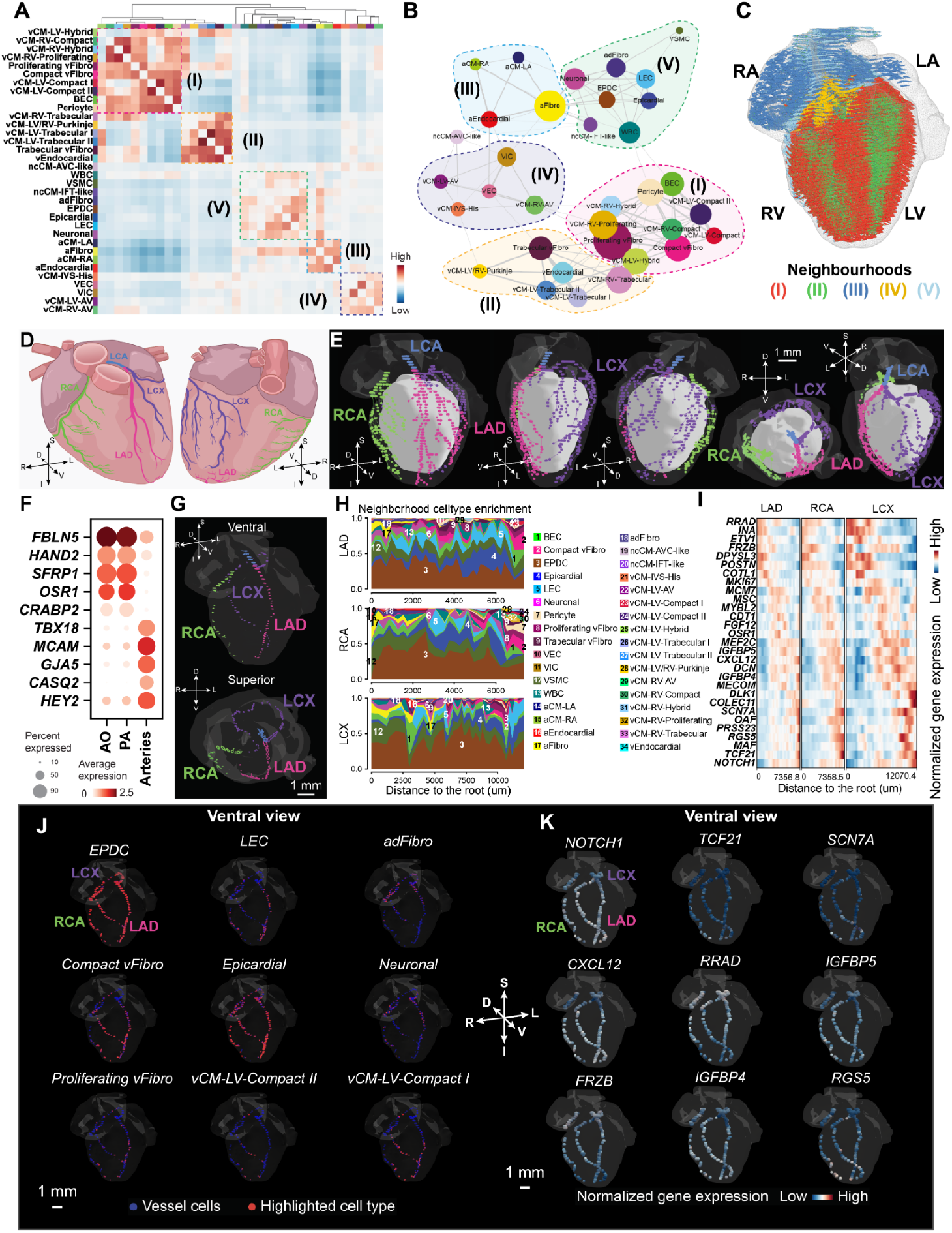
Spatial neighborhoods in the 3D developing heart. (**A**) Heatmap showing cell-type proximity scores across all 34 cell types defined in Figure 4F. Dendrogram on top marks the hierarchical clustering of the proximity scores across cell types. Five spatial neighborhoods are annotated using boxes. The scale bar in the bottom right indicates the strength of proximity score. (**B**) Proximity network graph illustrates positive associations among cell types (represented by different nodes in the graph), where the node size reflects the number of cells for that particular cell type and edge width indicates strength of association. Nodes are grouped into different spatial neighborhoods. (**C**) 3D spatial distribution of the cells comprising the five neighborhoods (colored as indicated) across a half bisected heart along the central left-right plane. (**D**) Schematic of the left-dominant human heart descending arteries drawn based on^101^. The aorta (AO), pulmonary artery (PA), left coronary artery (LCA), left anterior descending artery (LAD), left circumflex artery (LCX), and right coronary artery (RCA) are colored. (**E**) Multiviews of the 3D-reconstruction from Figure 4G onto which the corresponding arteries in (D) are colored (see **STAR Methods** for details). (**F**) Dot plot showing the expression profiles of highly differentially expressed genes in the aorta (AO), pulmonary artery (PA), and coronary arteries. (**G**) 3D spatial distribution of three long branches of the coronary arteries (LAD, LCX, and RCA). (**H**) Stacked area plot illustrating the neighborhood cell type composition along three branches from root to distal tip. Each colored region represents a distinct cell type numbered and colored as in the legend on the right. (**I**) 3D spatial distributions of 9 representative cell types enriched in the neighborhood of the selected vessels, where red dots indicate cells from the enriched cell type. **(J)** Gene expression heatmaps along the LAD, RCA, and LCX branches from root (left on the heatmap) to distal tip (right on the heatmap). Each row represents a different gene, and the color scale indicates relative expression (blue for lower expression and red for higher). **(K)** 3D expression maps for 9 representative genes in the selected vessels, where blue indicates lower expression and red - higher expression.

We observed that neighbourhood V contained multiple non-cardiomyocyte cell types including the epicardial cells, epicardium-derived cells (EPDCs), adventitial fibroblasts (adFibros), vascular smooth muscle cells (VSMCs), lymphatic endothelial cells (LECs) and neuronal-like cells (**Figures 5A-5C**). Many of these cell types are known to migrate during heart development from the surface of the heart towards the ventricular wall^49,51^. Among these cell types, we focused on VSMCs as landmarks to characterize the cellular composition and expression gradient along the coronary arteries.

The coronary arteries are key cardiac structures, whose defects cause major heart diseases^52–55^ yet whose characterization remains challenging using individual 2D sections. In contrast, the 3D heart reconstruction allowed us to trace the vascular smooth muscle cells (VSMCs expressing *MYH11*)^56^, and reconstruct the vasculature of the heart using a hierarchical tree-construction algorithm that links vessels cross sections (**STAR Methods**), enabling us to define the aorta (AO), the pulmonary artery (PA),left coronary artery (LCA), right coronary artery (RCA), and their major coronary artery segments (**Figures 5D and 5E)**. The LCA branched into the left circumflex artery (LCX) and the left anterior descending (LAD) artery (**Figures 5D and 5E**)^57^. The AO and PA VSMCs have differential gene expression compared to those of coronary arteries marked by elevated expression of *FBLN5, HAND2, SFRP1, OSR1*, and *CRABP2* (**Figure 5F**). These genes are known markers of neural crest cells (*CRABP2*)^58^ and second heart field (*HAND2, OSR1*)^59,60^ lineages, consistent with their developmental origin from the neural crest cells and second heart fiel^61,62^. In contrast, coronary artery VSMCs showed higher expression of *TBX18, MCAM, GJA5, CASQ2*, and *HEY2* (**Figure 5F**), reflecting the contribution of epicardium to VSMC as *TBX18* is the marker gene of epicardium^63^. Starting from the root region of the aorta, the left and right coronary arteries extended across the left and right ventricular surfaces, giving rise to a branching network that spans the entire length of the ventricles from base to apex, highlighting the ability of MERFISH+ to capture their trajectories in 3D (**Figures 5E and 5G**).

The 3D heart atlas allows for identifying the cellular environment and molecular pathways facilitating arterial morphogenesis. We quantified cell-type composition along the proximal–distal axis within the neighborhood of the VSMC layer (~2–3 cell diameters) for the longest branches of the RCA, LAD, and LCX^64^. Multiple cell types were proximal to arterial VSMCs, including epicardial cells, EPDCs, BECs, pericytes, fibroblasts, LECs and neural cells (**Figure 5H**), that are consistent with pro-angiogenic support by pericytes, EPDCs, BECs, and fibroblasts, and with artery co-development by neural cells and LECs^65–67 51,64^. Along the coronary long axis, several populations exhibited proximal–distal gradients (**Figures 5H and 5J**): compact ventricular cardiomyocytes (vCM-LV-Compact I/II; vCM-RV-Compact) and compact vFibro were enriched distally, aligning with the gradual integration of coronary vessels into the compact ventricular myocardium^64^. Epicardial cells were also enriched at distal segments, consistent with their capacity to differentiate into coronary smooth muscle cells ^63^.

In parallel to the cellular neighborhoods, we also quantified the gene expression in VSMCs along the proximal–distal axis of arterial branches (**Figures 5I and 5K**). Several genes exhibited differential expression along the proximal-distal axis, including *TCF21, CXCL12, NOTCH1, FRZB,RRAD* (**Figures 5I and 5K**). Higher expression at the distal ends of *TCF21*, a marker of epicardial-derived cells (EPDCs), may reflect EPDCs differentiating into VSMCs^68,69^. Analogously, higher expression at the distal ends of *NOTCH1* may reflect epicardial-to-VSMC differentiation in the coronary wall^70^. CXCL12, which has recently been identified as a key regulator of left and right coronary vessel branching during cardiac development ^26,71,72^, may be enriched at the distal ends to provide endothelial guidance cues for vessel formation ^26,71–73^. Together, these 3D analyses provide a spatially resolved catalog of transcriptional and cellular neighborhood changes in VSMCs along the developing coronary arteries, highlighting region-specific molecular programs that govern arterial maturation and elongation.

### Construction of a unified 3D multi-modal reference human heart atlas

We computationally integrated 2D transcriptomic data (**Figure 2**) and the spatially resolved histone modification maps (**Figure 3**) into a unified 3D reference (**Figure 6A left**). Building upon the variational autoencoder (VAE) framework^74^ for joint probabilistic modeling of multi-omic data, we developed Spateo-VI, a novel integration framework that explicitly models spatial proximity between cells based on the attention of a neighborhood graph to generate spatial multi-omics 3D profiles (**Figure 6A middle, STAR Methods**), enabling all downstream analyses (**Figures 6B-6I**). Specifically, gene expression profiles measured across 1,835 genes and the spatial histone modification data, originally acquired from 2D section, were mapped into the 3D heart transcriptome reference. This enhanced 3D multi-modal molecular hologram of the human developing heart opens doors to novel analyses, including the characterization of spatially enriched cell-cell interactions (**Figure 6A right**). Using this multi-modality integration, we extended COMMOT^75^ to implement COMMON-3D (COordinate-aware MultiMOdal Neighbor-based Cell-Cell interaction analysis in 3D space) to spatially infer interaction between specific ligand and receptor pairs with GPU acceleration (**STAR Method**).

**Figure 6:**
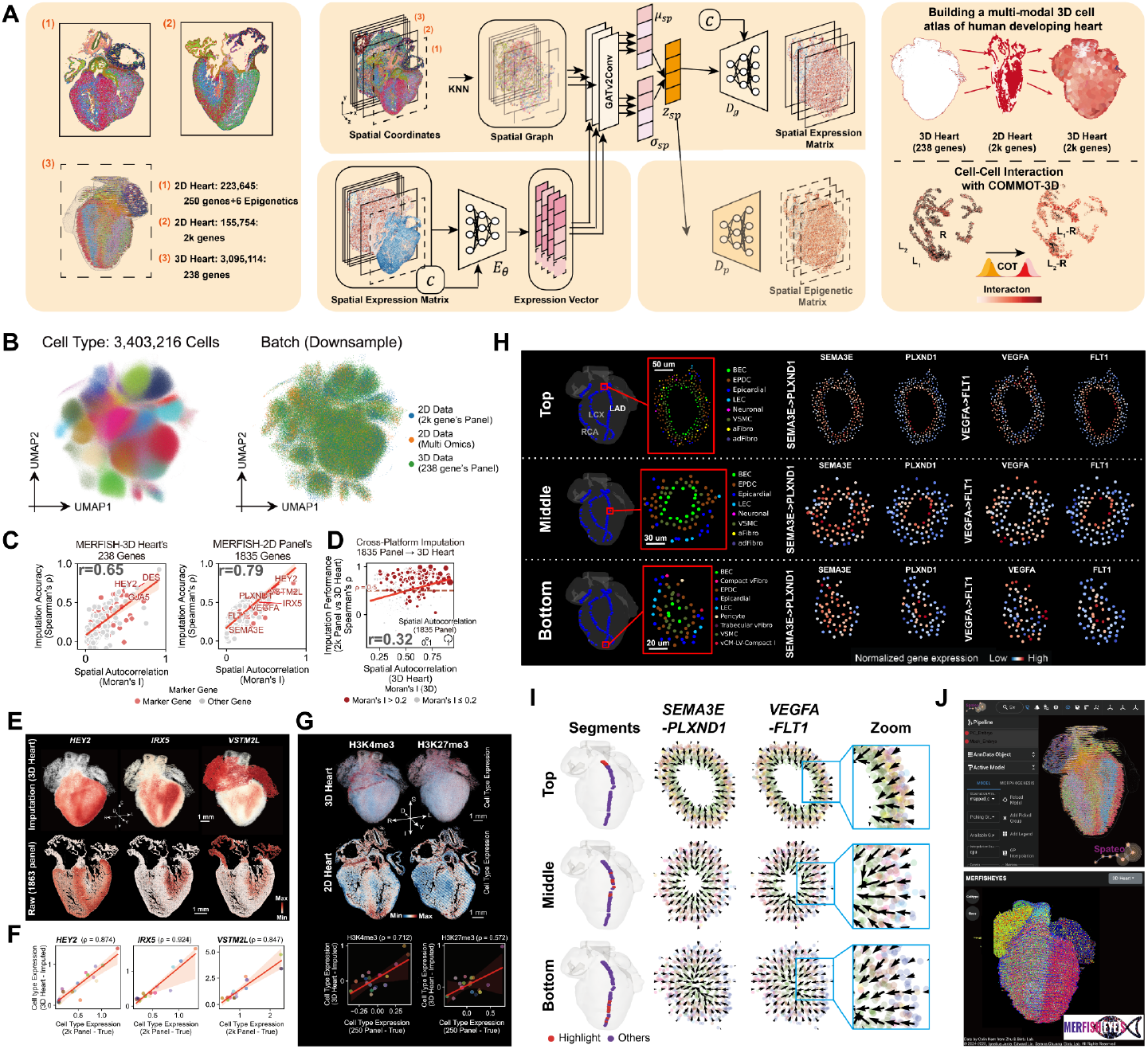
Computational integration of 2D spatial transcriptomic data and spatially resolved histone modification maps into a unified 3D reference. (**A**) Schematic of a new integration algorithm called Spateo-VI. Left panel: three different input data modalities; Middle panel: Schematic of the computational workflow integrating spatial coordinates, gene expression, and chromatin features using Spateo-VI. Right panel: Application of the model; top, projection of 2D multi-modal data (1,835-gene expression and epigenetic mark landscapes) into 3D heart space; bottom, inference of spatially enriched cell-cell interactions using COMMON-3D. (**B**) UMAP of the model’s latent space showing separation of transcriptionally distinct cell types across 3.4 million single cells, colored by cell types (left) or data input (right). (**C**) Relationship between imputation accuracy and spatial specificity. The y-axis represents imputation accuracy, measured by the correlation between true gene expression and imputed gene expression. The x-axis represents spatial specificity, measured by Moran’s I. The left plot shows analyses for the 238 genes measured in 3D heart, and the right plot shows analyses for the 1,835 genes measured in 2D heart. (**D**) Cross 3D-2D’s spatial specificity of imputation results at cell type level. The 1,835 genes are imputed for the 3D dataset and the average gene expression for each cell type is compared to that of the 2D MERFISH+ 1,835-gene panel experiment to calculate the Spearman’s ρ, similar as in panel F. (**E**) Spatial expression maps of representative imputed genes included *HEY2, IRX5* and *VSTM2L*. Top panel: the imputed expression in 3D heart. Bottom panel: the measured expression in 2D heart. (**F**) Correlation across cell types of the mean expression of three characteristic genes measured by MERFISH+ in 2D and the model-imputed data into the 3D reference. (**G**) Top: Imputed H3K4me3 and H3K27me3 histone mark distributions in 3D. Middle: The measured 2D histone mark distribution shown for comparison. Bottom: same as (F) but for H3k4me3 or H3K27Me3 respectively (r = 0.742 and r = 0.572). (**H**) Cross-sections of the LAD along 3 positions from root to distal tip showing cell types and imputed gene expression of two known ligand-receptor pairs *SEMA3E*-*PLXND1* and *VEGFA*-*FLT1*. (**I**) Spatial gradient of ligand-receptor interaction in cross-sections of the LAD. Three segments along the LAD from the root to the distal tip are projected to create an aggregate representation of the cross-section. Vector fields (represented by quivers) show the direction and magnitude of *SEMA3E*-*PLXND1* and *VEGFA*–*FLT1* signaling flow in the LAD. (**J**) Web portals for data access and exploration: Spateo-viewer (left) for 3D visualization and spatial analysis and MERFISHEYE (right) for 2D and 3D visualization.

To evaluate the performance of our model, we first visualized the latent space using UMAP and observed separation across 32 transcriptionally distinct cell types with minimal batch effect (**Figure 6B**). We found that imputation accuracy (Spearman rank correlation between predicted gene expression and the input ground-truth expression) correlated strongly with differential spatial expression: genes with higher Moran’s I spatial autocorrelation exhibited better imputation performance (**Figure 6C Left**, 238 genes in 3D heart**; Right**, 1,835 genes in 2D). We also evaluated the imputation accuracy in the 3D heart at the cell type level and found that genes’ imputation accuracy (based on cell-type correlation with 2D heart data) increased with higher Moran’s I, validating the model’s spatial sensitivity (**Figure 6D**). Spatial gene expression maps and correlation across cell types for representative genes, including *HEY2, IRX5*, and *VSTM2L*, demonstrated high concordance between imputed 3D patterns and experimental measurements from 2D sections (**Figures 6E and 6F**). Similarly, we validated the quality of the imputed chromatin modification signals, including H3K4me3 and H3K27me3 histone mark distributions in 3D (**Figure 6G top panels**) and found a high correlation across cell types with that of the measured 2D data (**Figure 6G bottom**), with Spearman’s correlation coefficients of ρ = 0.742 and ρ = 0.572, respectively.

Building on the 3D imputed transcriptional profiles, we investigated specific ligand receptor pairs, *SEMA3E*-*PLXND1* and *VEGFA*-*FLT1*, known to interact within the vascular niche in the developing embryo^76–79^. Within cross-sections of the left circumflex (LCX), left anterior descending (LAD), and right coronary artery (RCA) along the main branches, blood endothelial cells (BECs) constitute the innermost layer of the intima, forming a cellular lining of the vessel lumen (**Figures 6H-left, S9A and S9B)**. This layer is enclosed by vascular smooth muscle cells (VSMCs), which compose the medial layer of the artery wall, followed by EPDCs and other cell types in the outermost layer. The ligands *SEMA3E* and *VEGFA* show enrichment in VSMCs, whereas their receptors, *PLXND1* and *FLT1*, exhibit higher expression in BECs (**Figures 6H-right, S9A and S9B)**. This indicates a spatially coordinated pattern of ligand–receptor signaling relevant to coronary vessel function ^79^ and structure. To further examine the interaction flows of these pathways in three dimensions, we generated vector field representations illustrating the directionality and magnitude of *SEMA3E-PLXND1* and *VEGFA–FLT1*-mediated signaling using COMMON-3D (**Figures 6I and S9C**). This flow is consistent with the anatomical organization of the vessel wall and existing literature^76–79^, where the endothelium resides on the inner side and receives SEMA3E and VEGFA signals along the length of the blood vessels from surrounding cells, including vascular smooth muscle cells and other neighboring cell types.

To facilitate data access and community exploration, we developed two interactive web-based tools: Spateo-viewer, for 3D spatial transcriptomics visualization and analysis, and cell-cell interaction maps^26^ (https://viewer.spateo.aristoteleo.com/), and MERFISHEYES (https://zhu.merfisheyes.com/) for both 2D and 3D visualizations (**Figure 6J**).

In summary, this new computational framework, Spateo-VI, which integrates various 2D, and 3D spatial datasets, is developed to create the first 3D multi-modal map of human heart development. Furthermore, it offers a powerful approach for exploring and analysing single cell transcriptomic and epigenetic states and ligand-receptor interactions of cardiac development in 3D.

## Discussion

This study integrates three major methodological and data-driven advances that together establish a next-generation spatial genomics platform: 1) a molecular anchoring strategy which, together with new microscopy and microfluidics instrumentation, extends the capabilities of MERFISH into its next high-throughout form, MERFISH+; 2) a resource capturing the first molecularly defined 3D atlas of the developing human heart; and 3) novel computational tools that enable both the cellular and signaling characterization of intricate 3D cardiac structures and the multimodal imputations from 2D spatial imaging data onto a reconstructed 3D heart model.

### Advancing MERFISH through molecular anchoring and instrumentation

We characterized a probe-anchoring protocol in which spatial transcriptomic probes are covalently linked to polyacrylamide hydrogels cast on top of the tissue samples. This approach markedly enhanced sample stability and imaging longevity in MERFISH experiments. Unlike previous methods for increasing probe stability that focused on anchoring the targeted RNA^80^ or enzymatically converting the targeted RNA into cDNA^81 82^, here we provide a cost-effective and enzymatic-free solution of anchoring the target hybridization probes rather than their substrate. We demonstrated that this method of anchoring is highly efficient (capturing more than 50% of the initial signal - **Figure 1**) and enabled stringent wash conditions between hybridization cycles that allows imaging experiments to be “reset” if needed, a particularly valuable feature when working with limited human samples. Moreover, this anchoring chemistry supported multi-month continuous imaging with consistent signal performance, enabling more than a hundred hybridization and imaging cycles. Here, we demonstrated this advantage progressively by first imaging over 1,800 genes in a modular fashion across 112 cycles of imaging (**Figure 2**) and then performing multi-modal imaging across 93 cycles of imaging capturing: the 50-nm scale of chromatin organization across tens of genomic loci, the nascent RNA and mature RNA across hundreds of genes and the nuclear accumulation of epigenetic marks via iterative immunofluorescence (**Figure 3**). Finally, with RNA stability and imaging time no longer limiting factors, we re-engineered the microfluidics and microscopy system to enable 10 fold larger imaging area compared to our prior transcriptomic and multimodal imaging studies^15 40^ and 4-fold larger than what is commercially available for MERFISH or other imaging-based spatial transcriptomics platforms. This complete imaging platform we call MERFISH+ enabled the molecular profiling of more than 1.5 million cells per experiment at a cost of less than a thousand dollars in total reagent cost.

### Building the first 3D molecular atlas of the human heart

Leveraging these advances, we imaged an intact human developing heart with two large-scale MERFISH+ experiments. Empowered with 3D molecular alignment tools (Spateo) we reconstructed a comprehensive 3D map of the developing human heart at whole-organ scale. The resulting dataset contained 3.1 million cells across 34 distinct populations, including under-characterized cell types localized to specialized cardiac structures, such as valves and the vascular system (**Figures 4 and 5**). Using novel variational autoencoders designed specifically to take advantage of both transcriptional and spatial cues we imputed the measurements of 1,835 gene transcripts and the multimodal epigenetic marks into the 3D heart model. This resource provides a comprehensive molecular and anatomical framework for understanding how cellular populations assemble into large scale anatomical structures. We have made this data available and explorable via our visualization tools MERFISHEYES and Spateo-viewer. The reconstructed 3D atlas revealed molecular correlates of structural heterogeneity. Quantitative analysis of ventricular wall organization identified distinct contributions of trabecular, hybrid, and compact cardiomyocytes to local wall thickness (**Figure 4**). Similarly, 3D reconstructions of coronary arteries revealed their cellular compositions and transcriptional heterogeneity, which are features difficult to resolve from 2D data. Along the proximal-distal axis of coronary vessels, we detected distinct spatial domains of endothelial and mural cell subtypes accompanied by ligand-receptor signaling gradients (e.g., SEMA3E and VEGFA), highlighting potential mechanisms guiding vascular development.

### Broader implications and outlook

Beyond the cardiac focus within this manuscript, we believe that the MERFISH+ platform and its computational frameworks establish a generalizable and scalable approach for large format spatial genomics. The same strategy can be extended to construct multimodal, spatially resolved cell atlases of other complex human organs, including the brain, kidney, and lung, where existing efforts have been largely limited to 2D. By combining rich volumetric datasets with emerging machine learning tools, including generative AI models for 3D tissue synthesis and disease simulation, MERFISH+ provides a foundation for building virtual organs that bridge molecular and anatomical understanding. We anticipate that these integrated advances will accelerate both basic biological discovery and translational modeling across human organ systems.

## Acknowledgements

This work was in part supported by the National Institute of Mental Health (DP5-OD031878 to B.B.), 1S10OD032391-01 to QZ, CZI AICP-0000000175 to BB and QZ, Sutarjd family donation to XQ, YL and ZZ, Arc Impetus Award to XQ, YL. Illustrations in Figures 1A, 4C, and Supplemental Figures 1 and 2 were created using BioRender.

## Author contributions

Colin Kern, designed the MERFISH+ imaging experiments, developed and optimized MERFISH+ imaging data analysis pipelines, analyzed MERFISH+ imaging data, wrote the manuscript, designed and generated figures.

Qingquan Zhang; conceived and designed the heart studies, performed sample preparation and experiments, annotated and organized the data, results, and figures, and wrote the manuscript.

Yifan Lu; developed spatial computational analysis methods for reconstructing and analyzing the 3D heart reference, wrote the manuscript, designed and generated figures.

Jacqueline Eschbach; analyzed MERFISH+ imaging data, annotated descending arteries, wrote the manuscript, designed and generated figures.

Elie N Farah; Analyzed and annotated the data, helped provide biological interpretation/insights

Chu-Yi Tai:: Hybridized the heart samples and assisted with imaging of the MERFISH+ experiments

Kaifu Yang:: Analyzed MERFISH+ and Hi-C data, wrote the manuscript

Zoey Zhao:: assisted MERFISH sample processing and data analyses

Carlos Garcia Padilla:: analyzed the multiplexed DNA imaging data

Zehua Zeng:: developed Spateo-VI to integrate the 2D spatial omics data into the 3D reference.

Alexander Monell, contribution to probe design of MERFISH+ technology

Siavash Moghadami, contribution to probe design of MERFISH+ technology

Fugui Zhu, performed epigenetic marker antibody staining experiments

Yao, Fenyong, tissue handling and sample collection

Bin Li, assisted MERFISH data storage

Angie Hou, generated and curated MERFISH cell type lists, revised figures and manuscript.

Ignatius Jenie,: made the MERFISHEYES website.

Grant Tucker,: revised figures and manuscript

David Ellison, provided GPU computing resources for this project.

Neil Chi, supervised the study as well as conceived, designed and interpreted the heart studies and wrote the manuscript

Xiaojie Qiu:: conceived and designed the study of the data analyses of 3D heart atlas and wrote the manuscript.

Quan Zhu:: supervised the study, conceived and designed the MERFISH+ technology, conceived, designed and supervised the heart MERFISH+ imaging studies, and wrote the manuscript.

Bogdan Bintu:: supervised the study, conceived and designed the MERFISH+ technology, conceived, designed and supervised the heart MERFISH+ imaging studies, and wrote the manuscript.

## STAR★Methods

### RESOURCE AVAILABILITY

Further information and requests for resources and reagents should be directed to corresponding authors. Lead contact: Bogdan Bintu

## Materials availability

Templates and reagents for making these probes can be purchased from commercial sources, as detailed in the Key Resources Table.

## Code availability

- Analysis code used for MERFISH+ probe library design, image decoding, and data quantification and analyses is available at: https://github.com/deprekate/LibraryDesigner https://github.com/epigen-UCSD/MERlin https://github.com/epigen-UCSD/MERFISH_Plus_Paper https://github.com/aristoteleo/MERFISHVI https://github.com/aristoteleo/Human_Heart_MERFISH_Analysis_2024
- Code for controlling custom pipetting robot is available at: https://github.com/BogdanBintu/PipettingRobot

Any additional information required to reanalyze the data reported in this paper is available from the lead contact upon request.

### Key resources table

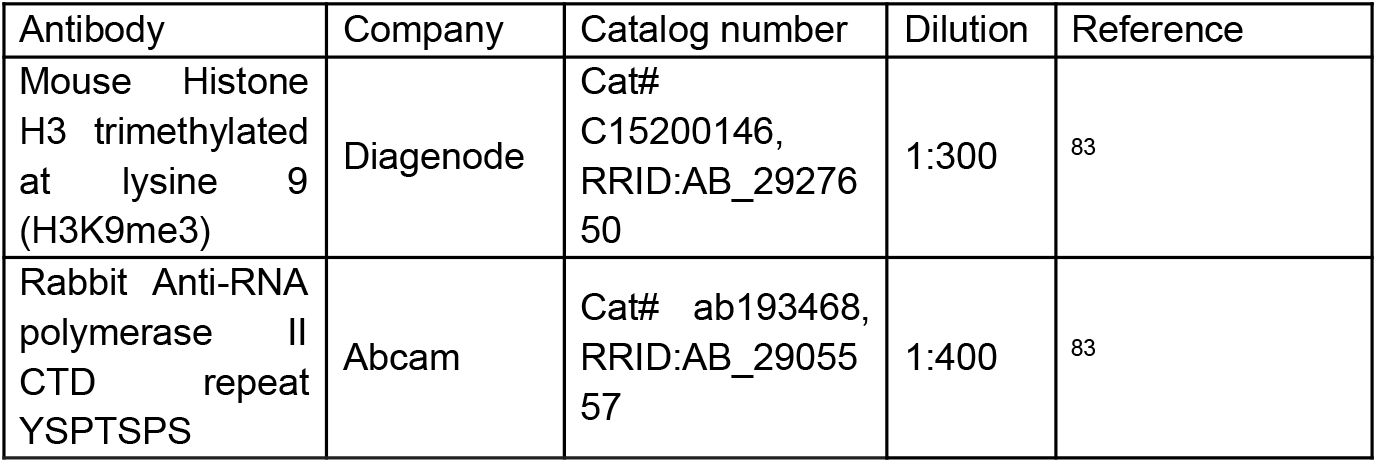
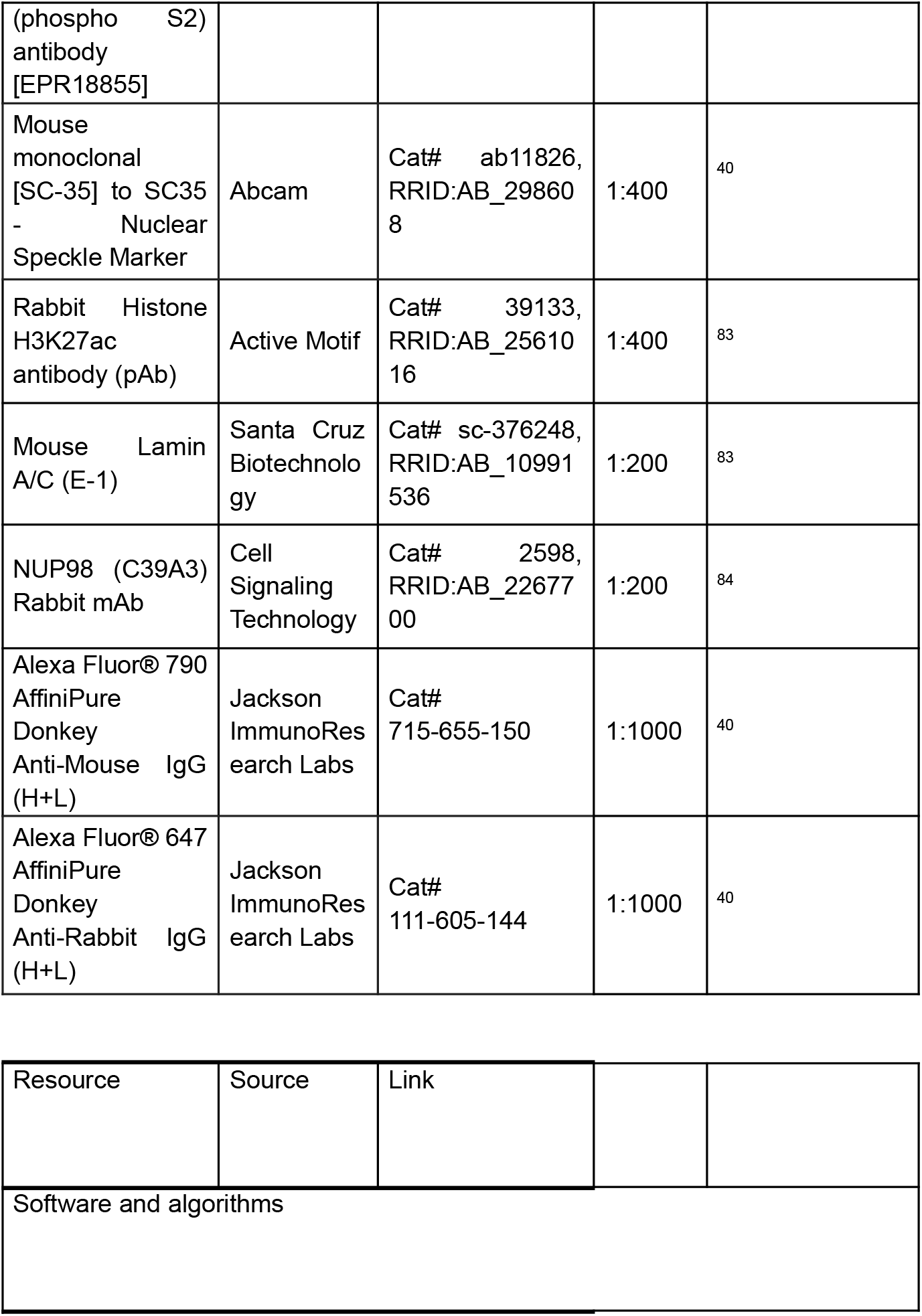
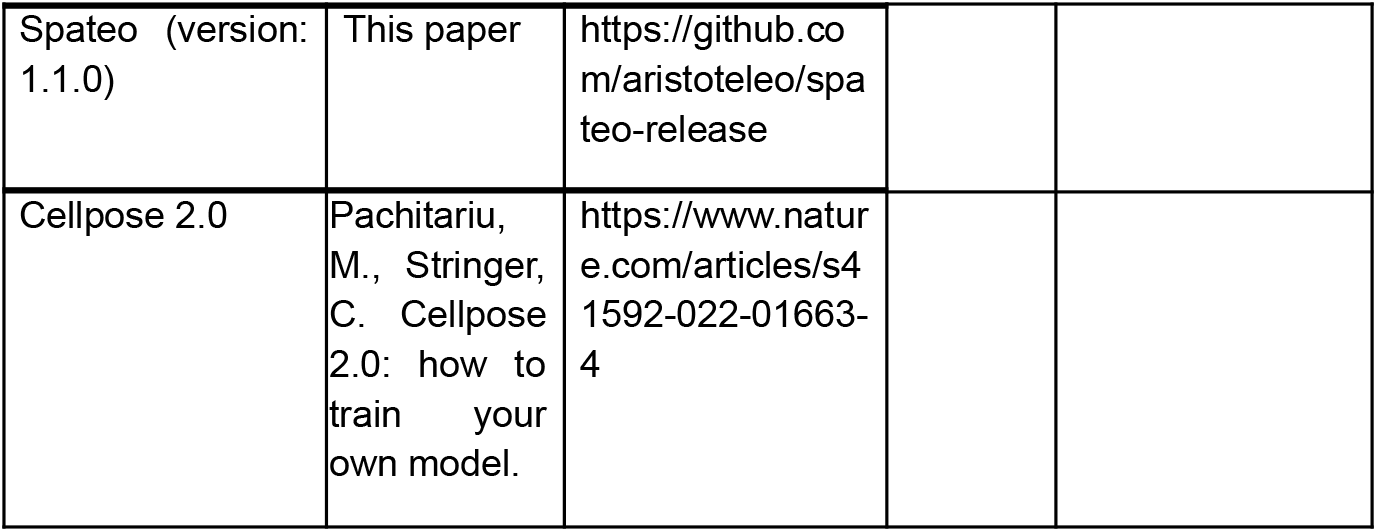

## Experimental model and study participant details

### Tissue samples

Heart samples were collected in strict observance of the legal and institutional ethical regulations. The heart samples were collected under a University of California San Diego (UCSD) Human Research Protections Program Committee Institutional Review Board (IRB)-approved protocol (IRB number 172111) by the UCSD Perinatal Biorepository’s Developmental Biology Resource after informed consent was obtained from the donor families. All experiments were performed within the guidelines and regulations set forth by the IRB (IRB number 101021, registered with the Developmental Biology Resource). Ethical requirements for data privacy include that sequence-level data be shared through controlled-access databases.

### Tissue processing

Tissue samples were dissected in buffer containing 125 mM NaCl, 2.5 mM KCl, 1mM MgCl_2_, and 1.25 mM NaH_2_PO_4_ under a stereotaxic dissection microscope (Leica). Samples for MERFISH were washed 1 time with ice-cold PBS, then fixed in 4% PFA at 4°C overnight. On the second day, the sample was washed in ice-cold PBS three times, 10 minutes each, and were incubated in 10% and 20% sucrose at 4°C for 4 hours each, and in 30% sucrose overnight, followed by immersion with OCT (Fisher, cat# 23-730-571) and 30% sucrose (1 vol:1 vol) for 1 hour. The sample was then embedded in OCT and stored at −80°C until sectioning.

A 12 p.c.w. human heart was systematically cryosectioned in the transverse plane at 16 μm thickness per slice, starting from the atria and progressing toward the apex. Each tissue section was mounted sequentially and serially onto a set of eight large-format glass slides in a round-robin fashion. Each individual slide therefore contained 21 sections, representing every 8th section along the atrial-to-apical axis. This strategy ensured that each of the eight slides captured a spatially even and parallel representation of the entire heart, preserving the full diversity of cell types, anatomical zones, and structural features across all slides. The process continued until atrial tissue was no longer present, at which point the superior portion of the heart was separately sectioned and similarly distributed. For this superior region, the remaining heart tissue was sectioned and again distributed across a second set of eight slides, each containing 32 tissue sections. This maintained the same 1-in-8 sampling strategy and ensured uniform representation of all anatomical regions across the second slide set.

The total number of sections is 53 x 8 = 424 sections with a mean thickness of 16 um, spanning a total length of 7 mm. One glass slide containing either 21 or 32 sections was selected, and the tissue was transferred onto a salinized coverslip. The coverslip was then mounted into the newly developed fluidics system for downstream analysis^12^. Subsequently, one coverglass was subjected to rapid fixation, stained with the same 238 genes^15^. Image processing and decoding was performed using a modified version of MERlin ^85^, which integrates Cellpose for cell segmentation and includes an alternative algorithm for drift correction that uses DAPI staining rather than fiducial beads and can correct for drift along the z-axis. We then performed a 3D reconstruction of the tissue sections using Spateo ^86^.

### Extended MERFISH+ probe library (1835 genes) design and construction

We developed custom primary probes using a Python-based workflow available at https://github.com/deprekate/LibraryDesigner. Probe sequences were selected following a previously established method^40^, in which each gene was targeted by a set of 5 to 60 unique 40-mer sequences complementary to its mRNA.

Probe selection involved:

1. Creating a k-mer index (17-mers) from the unspliced human transcriptome to asses sequence uniqueness
2. Screening candidate 40-mer regions for off-target potential based on their 17-mer content
3. Applying filtering criteria to retain only the most specific and effective sequences We then used transcript sequences from the human reference genome (hs1). Genes that were either too short or highly homologous to other targets were excluded, resulting in final panels of 238 and 1835 genes. To each 40-mer, we appended three unique 20-nt readout sequences per gene, along with universal PCR priming sites at both ends. All readout and primer sequences were checked against the human transcriptome to minimize cross-reactivity.

The probes were synthesized from custom oligo pools (Twist Biosciences), amplified via limited-cycle PCR, and purified using Zymo DNA oligo cleanup kits (D4003). RNA was transcribed in vitro using T7 polymerase (NEB), then reverse-transcribed into single-stranded DNA using Maxima reverse transcriptase (Thermo Scientific). RNA templates were removed through alkaline hydrolysis, followed by another round of DNA purification (Zymo Research, D4006). The forward primer used in PCR and reverse transcription carried a 5’-acrydite modification to enable covalent attachment of probes to the acrylamide hydrogels during tissue processing.

### Linker Probes and Codebook Construction

Each gene-specific linker probe consisted of a 20-nt region complementary to the gene’s readout sequence, followed by two tandem amplification binding sequences (each 20nt). Linkers were ordered from IDT in 384-well plates.

To assign spatial barcodes for MERFISH imaging, we designed a binary codebook using a custom Python script, available at https://github.com/epigen-UCSD/MERFISH_Plus_Paper. A binary code was randomly assigned to each gene such that each barcode contained exactly four 1’s across the chosen number of imaging rounds N, while maintaining a Hamming distance of four between codes to ensure error correction. We further refined the codebook using a Metropolis-Hastings optimization algorithm ^87^, minimizing the overlap of high-expression genes within the same hybridization round based on single-cell RNA-seq data. The final hybridization mixtures were prepared using a custom pipetting robot built by combining a CNC Router (CNC-4040-Basic, FoxAlien) with a programmable syringe pump from Hamilton. This robot dispensed gene-specific linkers into 1.5mL Lo-Bind tubes according to their assigned hybridization rounds. Code for operating this custom pipetting robot is provided at: https://github.com/BogdanBintu/PipettingRobot

### Amplification and Readout Probes

Amplification probes were designed in two tiers. Level-1 probes included a 20-nt arm to bind to the linker overhang, followed by four binding sites for the Level-2 probes. Level-2 probes were similarly designed to bind Level-1 probes and carry four additional sites for fluorescent readout probe binding. All amplification oligonucleotides were synthesized by IDT. Readout probes were 20-nt oligos, double-labeled with fluorescent dyes specific to the imaging channels: Cy3 (561nm), Cy5 (638nm), and Alexa Fluor 750 (750nm). These were purchased from IDT with high-performance liquid chromatography (HPLC) purification.

### Probe library design for RNA-MERFISH and chromatin tracing

We selected 40bp target sequences for DNA or RNA hybridization by considering each contiguous 40bp subsequence of each target of interest (the mRNA of a targeted gene or the genomic locus of interest) and then filtering out off-targets to the rest of the transcriptome/genome including repetitive regions, or too high/low GC content or melting temperature (TM). More specifically, our probe design algorithm was implemented with three steps: 1) Build a 17-mer index based on reference genome hs1 assembly (DNA) or the hg38 transcriptome (RNA). 2) Quantify 17-mer off-target counts for each candidate 40bp target sequence. 3) Filter and rank target sequences based on predefined selection criteria as previously described ^88,40^. The primary probe set was constructed for pan-tissue imaging (human brain, heart, lung, and pancreas) and included up to 60 unique 40-mer sequences per gene, selected to hybridize to either mRNA or genomic DNA. Target sequence design was carried out as previously described^40,88^, with modifications to ensure broad tissue compatibility. Target design proceeded through three main steps: (1) A 17-mer index was constructed from the human genome assembly hs1 for DNA targets and the hg38 transcriptome for RNA targets; (2) Each contiguous 40-mer candidate from the target mRNA or genomic locus was scanned for potential off-targets by quantifying 17-mer matches across the genome or transcriptome; and (3) candidate sequences were filtered and ranked based on predefined criteria, including melting temperature, GC content, repetitive element masking, and minimal off-target potential. Genes that failed to meet design criteria due to sequence redundancy or insufficient unique regions were excluded. Ultimately, 1,826 genes were selected for the final probe set. For each probe, additional sequence handles were appended, including three 20-nt gene-specific readout sequences and two 20-nt PCR primer sites. Readout sequences and primers were selected to minimize cross-hybridization, using prior design frameworks ^40^ and screened against the human transcriptome. Oligonucleotide pools (Twist Biosciences) were PCR-amplified using limited-cycle PCR, incorporating a 5′-acrydite modified forward primer to enable acrylamide gel incorporation. Amplified DNA was used for in vitro transcription using T7 polymerase (NEB), followed by reverse transcription with Maxima RT (Thermo Scientific) and RNA strand hydrolysis to yield single-stranded DNA probes, which were then purified (Zymo Research D4003 and D4006 kits).

### MERFISH+ gene selection

The gene panels used in MERFISH+ experiments were selected based on prior snRNAseq to allow for a high definition of cell types across multiple tissues. 51% of these selected genes also had high differential expression in snRNAseq of the heart ^15^ that are defined as being expressed in at least 50% of a celltype and have at least double the average expression level compared to any other cell type.

The MERFISH gene selection was performed by first using a scRNA-seq dataset from 12-15 p.c.w. heart^15^and using NSForest v2 ^89^ with default parameters to identify marker genes for the cell type clusters in this data. This list of genes was supplemented with additional marker genes from the literature. The target sequences for each gene were concatenated with one or two unique readout sequences to facilitate MERFISH or smFISH imaging.

### Design of chromatin probes

We designed probes for DNA hybridization similarly as those for the RNA MERFISH as described in ^40,90^. Briefly, we first partitioned FXYD7 locus into 10-kb segments. After screening against off-target binding, GC content and melting temperature, ~100 unique 40 bp target sequences were selected for each 10-kb segment. We concatenated a unique readout sequence to the target probes of each segment to facilitate sequential hybridization and imaging of each locus.

### Acrydite probe synthesis

The acrydite probes were generated using oligonucleotide pools, following the previously described method ^20^. Firstly, we amplified the oligopools (Twist Biosciences) using a limited cycle qPCR (approximately 15 – 20 cycles) with a concentration of 0.6 µM of each acrydite primer to create templates. These templates were converted into RNAs using the in vitro T7 transcription reaction (New England Biolabs, E2040S). These RNAs were then converted back to single-stranded DNA using reverse transcription with a concentration of 16 µM of each acrydite primer. Subsequently, the DNA oligos were purified using alkaline hydrolysis to remove RNA templates and cleaned with columns (Zymo Research, D4060). The resulting acrydite probes were stored at −20 °C.

### RNA MERFISH sample preparation

Fresh frozen hearts were sectioned at −20°C using a Leica CM3050S cryostat. Series coronal sections of 16 μm thickness were performed at ~600 μm along the anterior-posterior axis of human hearts to capture the major cardiac structures. Sections were collected on salinized and poly-L-lysine (Millipore, 2913997) treated 40-mm, round no.1.5 coverslips (Bioptechs, 0420-0323-2). After drying for 20 min, tissue sections were stored at −80°C until use.

MERFISH measurements of 238 genes with 200 non-targeting blank controls was performed as previously described, using the published encoding sequences and readout probes^15^, subsequently pre-cleared by immersing into 30% (vol/vol) ethanol, 50% (vol/vol) ethanol, 70% (vol/vol) ethanol, and 100% ethanol, air dried for 5 minutes, then, 70% (vol/vol) ethanol, 50% (vol/vol) ethanol, each for 5 minutes. The tissue was then treated with Protease III (ACDBio) at 40 °C for 15 minutes prior to being washed with 1x PBS for 5 minutes at room temperature. Then the tissues were preincubated with hybridization wash buffer (40% (vol/vol) formamide in 2x SSC) for ten minutes at room temperature. After preincubation, the coverslip was moved to a fresh 60 mm petri dish and residual hybridization wash buffer was removed with a Kimwipe lab tissue. In the new dish, 50 μL of encoding probe hybridization buffer (2X SSC, 50% (vol/vol) formamide, 10% (vol/vol) dextran sulfate, and a total concentration of 5 uM encoding probes and 1 μM of anchor probe: a 15-nt sequence of alternating dT and thymidine-locked nucleic acid (dT+) with a 5′-acrydite modification (Integrated DNA Technologies). The sample was placed in a humidified 47^°^C oven for 18 to 24 hours then washed with 40% (vol/vol) formamide in 2X SSC, 0.5% Tween 20 for 30 minutes at room temperature. Samples were post-fixed with 4% (vol/vol) paraformaldehyde in 2X SSC and washed with 2X SSC with murine RNase inhibitor for five minutes. To anchor the RNAs in place, the encoding-probe–hybridized samples were embedded in thin, 4% poly-acrylamide (PA) gels as described^15^. Briefly, the hybridized samples on coverslips were washed with a de-gassed 4% poly-acrylamide solution, consisting of 4% (vol/vol) of 19:1 acrylamide/bis-acrylamide (BioRad, 1610144), 60 mM Tris⋅HCl pH 8 (ThermoFisher, AM9856), 0.3 M NaCl (ThermoFisher, AM9759) supplemented with the polymerizing agents ammonium persulfate (Sigma, A3678) and TEMED (Sigma, T9281) at final concentrations of 0.01% (wt/vol) and 0.05% (vol/vol), respectively, for 10min to equalibrate. Then the sample was reverted onto a glass plate that is cleaned and siliconized and spoted with 300ul of the 4% (vol/vol) of 19:1 acrylamide/bis-acrylamide (BioRad, 1610144), 60 mM Tris⋅HCl pH 8 (ThermoFisher, AM9856), 0.3 M NaCl (ThermoFisher, AM9759) supplemented with the polymerizing agents ammonium persulfate (Sigma, A3678) and TEMED (Sigma, T9281) at final concentrations of 0.03% (wt/vol) and 0.15% (vol/vol), respectively. The gel was then allowed to cast for 5 h at room temperature. The coverslip and the glass plate were then gently separated, and postfixed with 4% PFA, then the PA film was incubated with a digestion buffer consisting of 50 mM Tris⋅HCl pH 8, 1 mM EDTA, and 0.5% (vol/vol) Triton X-100 in nuclease-free water and 1% (vol/vol) proteinase K (New England Biolabs, P8107S). The sample was digested in the digestion buffer for ~5 h in a humidified, 37°C incubator and then washed with 2xSSC three times. MERFISH measurements were conducted on a home-built system as described^5^ and below.

### Multi-modal chromatin tracing sample preparation

The sample preparation is similar to that for RNA MERFISH sample preparation with the exception that the sample is treated with 0.5% Triton X-100 in 1X PBS for 10 minutes at room temperature instead of ethanol dehydration. Briefly, Frozen hearts were sectioned at −20°C using a Leica CM3050S cryostat. Series coronal sections of 14 μm thickness. Sections were collected on salinized and poly-L-lysine (Millipore, 2913997) treated 40-mm, round no.1.5 coverslips (Bioptechs, 0420-0323-2). After drying for 20 min, Fixed tissue sections were permeabilized with 0.5% Triton X-100 (Sigma-Aldrich, T8787) in 1× PBS with RNase inhibitors for 10 min at room temperature and washed once with 1X PBS with RNase inhibitors for 3 min. After permeabilization, coverslips were moved to a fresh 60 mm petri dish and treated with 0.1M hydrochloric acid (Thermo Scientific™, 24308) for exactly 5 min at room temperature, and washed three times for 5 min with 1× PBS with RNase inhibitors. Tissue sections were preincubated with the hybridization wash buffer (50% (vol/vol) formamide (Ambion, AM9342) in 2x SSC (Corning, 46-020-CM)) for 10 min at room temperature. Prepared a layer of fresh parafilm with a 0.8 cm diameter hole in the center and placed in a fresh 60 mm petri dish. Then 50 uL of encoding probe hybridization buffer (50% (vol/vol) formamide (Ambion, AM9342), 2x SSC (Corning, 46-020-CM), 10% Dextran Sulfate (Millipore, S4030), 5 μM encoding probes and 1 μM of anchor probe, a 15-nt sequence of alternating dT and thymidine-locked nucleic acid (dT+) with a 5′-acrydite modification (Integrated DNA Technologies) was added to the center. Coverslips were wiped away excess hybridization wash buffer with Kimwipes and placed tissue-side-down onto the droplet. Samples were in a humidity chamber at 47 °C for 18 - 24 hours. The sample was then washed with 40% (vol/vol) formamide in 2X SSC, 0.5% Tween 20 for 30 minutes and post-fixed with 4% (vol/vol) paraformaldehyde in 2X SSC followed by two washes in 2X SSC with murine RNase inhibitor for 5 min each at room temperature before being embedded in thin, 4% polyacrylamide (PA) gels as described for RNA MERFISH sample ^21^.

### Microscopy and fluidics equipment set up

#### 1) Microscope Setup for Image Acquisition

Image acquisition was carried out using a custom-built microscope system, as previously described ^12,40,91^. The core of the setup was a Nikon Ti-U microscope body fitted with a Nikon CFI Plan Apo Lambda 60x oil immersion objective (NA 1.4).

Illumination was provided based on: Light Engine Illumination (Lumencor). Specifically, A Lumencor CELESTA light engine (wavelengths: 405, 477, 546, 638, 749 nm) was used with a penta-bandpass dichroic (IDEX, FF421/491/567/659/776-Di01) and filter (IDEX, FF01–441/511/593/684/817). In most experiments, beam uniformity was enhanced using either a refractive beam shaper (Newport Optics, GBS-AR14) or a vibrational optical fiber (Errol, custom Albedo unit). Images were acquired with a Kinetix 10 camera (Teledyne Action Optics). Sample positioning was controlled with a motorized XYZ stage (Ludl), and focus was maintained using a custom-built autofocus system ^92^, employing IR lasers (Thorlabs LP980-SF15) and a CMOS camera (Thorlabs, uc480). All hardware components were synchronized and operated via a DigiKey DAQ card (DigiKey 781048-01) and a custom software.

#### 2) Fluidics System

The fluidics system consisted of a syringe pump (Gilson MINIPLUS 3), a 24-port Vici valve (VICI C25G24524UMTA), a flow chamber (Bioptechs 060319–2), and tubing sealed with pressure adhesive (Blu-tack). Each valve was connected to a distinct buffer, with dedicated lines for imaging, stripping, and washing. Buffers were delivered to the sample chamber, and waste was collected downstream in an open-loop setup.

The system supported hundreds of hybridization rounds, depending on buffer configuration. For experiments exceeding this capacity, buffer replacement was performed by:

1. Disconnecting the chamber and flushing the valve lines with 50% formamide and water, followed by 2xSSC.
2. Introducing new buffers.
3. Reconnecting the chamber and resuming imaging.

#### 3) Experimental Control Software

All components were controlled via custom software available at https://github.com/ZhuangLab, composed of the following modules:

1. Hal: Controls illumination, microscope components, and imaging parameters.
2. Steve: Acquires mosaic images and selects imaging regions.
3. Kilroy: Controls fluidics and executes buffer exchange sequences.
4. Dave: Automates the experiment by integrating Hal and Kilroy operations.

At experiment start, imaging parameters and buffer sequences were configured in Hal and Kilroy. A DAPI mosaic was acquired using Steve to define imaging regions. The full experiment protocol was loaded into Dave for automated execution. For long experiments, buffer exchanges were manually performed between Dave runs as needed.

The CNC liquid handler consists the following parts

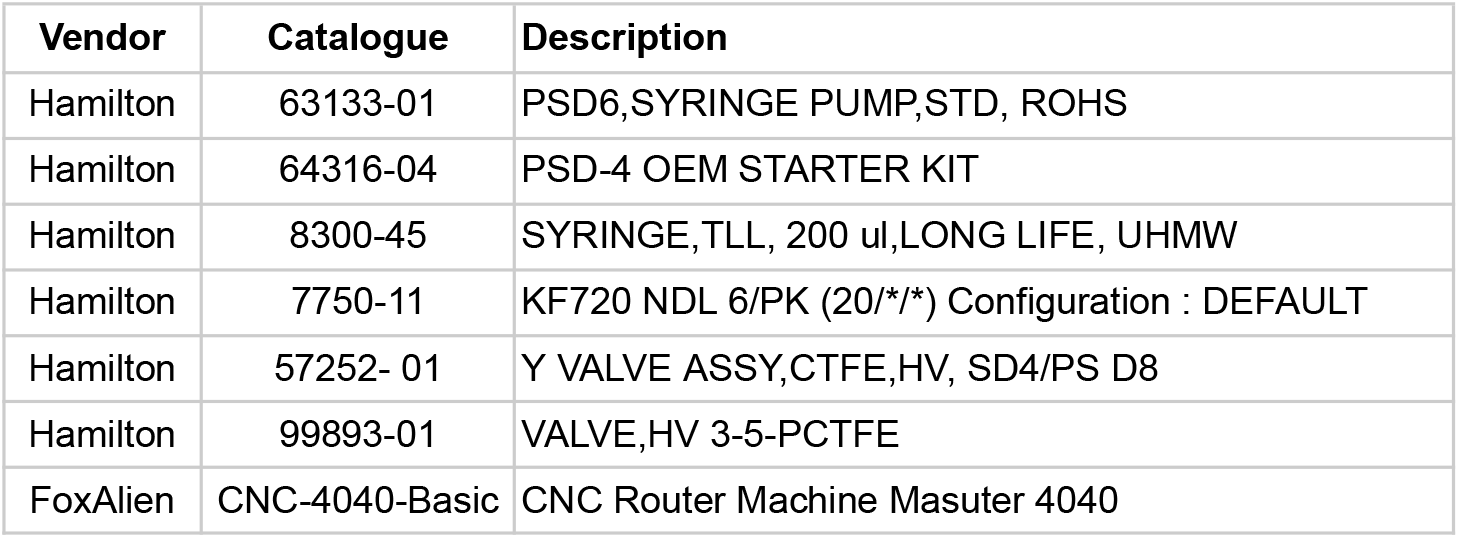

| Vendor | Catalogue | Description |
| --- | --- | --- |
| Hamilton | 63133-01 | PSD6,SYRINGE PUMP,STD, ROHS |
| Hamilton | 64316-04 | PSD-4 OEM STARTER KIT |
| Hamilton | 8300-45 | SYRINGE,TLL, 200 μl, LONG LIFE, UHMW |
| Hamilton | 7750-11 | KF720 NDL 6/PK (20/*/*) Configuration : DEFAULT |
| Hamilton | 57252- 01 | Y VALVE ASSY,CTFE,HV, SD4/PS D8 |
| Hamilton | 99893-01 | VALVE,HV 3-5-PCTFE |
| FoxAlien | CNC-4040-Basic | CNC Router Machine Masuter 4040 |

### Imaging and adaptor/readout hybridization protocol

MERFISH measurements were conducted on a custom microfluidics-microscope system with the configuration previously described^40^. Briefly, the system was built around a high-speed and high-sensitivity camera (the Kinetix model from Teledyne) capable of imaging twice the area of the prior model of Hamamatsu CMOS cameras. A newer line of Nikon 60x oil immersion objective (1.4 NA) was also used for better spherical and chromatic aberration corrections. The different components were synchronized and controlled using a National Instruments Data Acquisition card (NI PCIe-6353) and custom software ^40^.

To enable multi-modal imaging we first sequentially hybridized fluorescent readout probes and then imaged the targeted genomic loci and then the targeted mRNAs. Then we performed a series of antibody stains and imaging. Specifically the following protocol was used in order:

1. 52 rounds of hybridization and chromatin tracing imaging, sequentially targeting the 21 loci using 1-color imaging ^40^
2. 16 rounds of hybridization and MERFISH imaging, combinatorially targeting 298 genes using 3-color imaging
3. 21 rounds of hybridization and smFISH imaging, targeting 42 introns using 2-color imaging
4. 4 rounds of sequential staining for 6 different antibodies using 1-color or 2-color imaging.

The protocol for each hybridization included the following steps:

1. Incubate the sample with adaptor probes for 30 minutes for DNA imaging or 75 minutes for RNA imaging at room temperature.
2. Flow wash buffer and incubate for 7 minutes
3. Incubate fluorescence readout probes (one for each color) for 30 minutes at room temperature.
4. Flow wash buffer and incubate for 7 minutes.
5. Flow imaging buffer. The imaging buffer was prepared as described previously ^40^ and additionally included 2.5μg/mL of DAPI.

Following each hybridization the sample was imaged and then the signal was removed by flowing 100% formamide for 20 minutes and then re-equilibrating to 2XSSC for 10 minutes.

RNA-MERFISH measurement of the 1,835 gene panel was performed with a modular of encoding scheme of 48-bit binary barcode and a Hamming weight of 4 (HW4). Therefore, in each hybridization round, ~75 adaptor probes were pooled together to target a unique subset of the 1,835 genes.

### Immunofluorescence staining

Antibody imaging was performed immediately after completing the DNA and RNA imaging. The sample was first stained for 4 hours at room temperature using 2 primary antibodies of two different species (mouse and rabbit), washed in 2XSSC for 15 minutes and then stained for 2 hours using 2 secondary antibodies for each target species conjugated with fluorescent dyes. Image acquisition: After each round of hybridization, we acquired z-stack images of each FOV in 4 colors: 750 nm, 647nm, 560 nm and 405 nm. Consecutive z sections were separated by 300 nm and covered 15 um of the sample. Images were acquired at a rate of 20 Hz.

## DATA ANALYSIS

### Analysis of 2D heart sections with acrydite-modified MERFISH probes

MERFISH data imaged using acrydite-modified MERFISH probes were analyzed using MERlin v0.6.1 exactly as previously described^15^. Cell types were assigned and UMAPs generated using the “ingest” function of the Scanpy tool to map the newly generated data onto the previously published MERFISH data ^93^.

### Analysis of 2D heart sections with MERFISH+ probes

MERFISH data in **Figures, 1, 2** and **3** were processed with custom scripts developed for MERFISH+ data.

#### Image Pre-processing

##### Flat-field correction

Images were first corrected to remove variations in illumination present on the edges and corners of images (flat-field correction) by calculating the per-pixel median brightness across the first round of imaging, separately for each color. Each image was then inversely scaled by this median brightness.

##### Deconvolution

Next, the images were deconvoluted using the Wiener algorithm with a custom point spread function (PSF) calculated for the specific microscope used for imaging to reduce noise and enhance the discreteness of fluorescent spots. Finally, a high-pass filter was used with a blur sigma of 30 to further improve signal-to-noise ratio.

##### Localization of Fluorescent Spots

After pre-processing images, local maxima within a 1-pixel radius were identified and further filtered to enforce a brightness of at least 3600 and a correlation with the point spread function of 0.25. The remaining local maxima were used for decoding.

##### smFISH quantification

For quantification of transcripts imaged individually as smFISH targets, the fluorescent spots identified in the previous step were further filtered based on a minimum brightness relative to their local background. This threshold was selected separately for each color channel, and manually tuned to remove noisy and dim fluorescence while keeping bright puncta.

##### MERFISH decoding

To resolve RNA molecules from the identified fluorescent spots across imaging rounds, spots were first clustered into groups across images within a range of 2 pixels after correcting for drift using the same method described above (MERlin). Due to the MERFISH codebook designed with a hamming weight of 4, clusters with at least 3 spots were kept to allow for single-bit-error correction, while clusters with 1 or 2 spots were discarded. For each cluster, a brightness vector was constructed using the spot brightness in each round, then L2 normalized and compared with the codebook similar to MERlin described above.

To filter noise from the resulting decoded transcripts, a score was calculated for each transcript based on three metrics: 1) the average brightness across the fluorescent spots, 2) the distance of the brightness vector to the nearest spot in the MERFISH codebook, and 3) the average distance of each fluorescent spot to the median of all spots. These metrics were combined into a single score by calculating the combined Fisher p-value against the distribution of each metric across all decoded transcripts. The distribution of scores for decoded transcripts with a blank barcode identity were then compared to the distribution of scores for decoded transcripts with a gene identity, and a score threshold was chosen which separates the peaks of each distribution.

##### Cell Segmentation

Cell boundaries were determined in 3 dimensions (3D) using Cellpose v2^94^. First, for each z-slice image, a 2D segmentation mask was generated using the DAPI channel with Cellpose’s “dapi” model, after flat field correcting and deconvoluting the DAPI image as described in “Image pre-processing”. Cells were then linked between adjacent z-slices to create a 3D segmentation.

##### Assigning transcripts to cells

Transcripts identified by both smFISH and MERFISH were assigned to cells by first adjusting the coordinates of each molecule by the calculated drift between the images used for segmentation and those in which the RNA were imaged. Next, the transcript coordinates were rounded to the nearest integer and assigned to the cell based on the value of the segmentation mask at each transcript’s coordinates.

### Analysis of 3D heart reconstruction

We firstly preprocessed the 3D heart data using MERlin as described in Farah et al., 2024^15^, and then reconstructed the 3D heart map using Spateo ^26^, with the following minor modifications for each method:

#### Localization of RNA molecules

MERFISH images generated for the 3D heart reconstruction were analyzed using a modified version of MERlin v0.6.1 to decode individual RNA molecules. Briefly, these modifications added support for images in compressed Zarr format and extended the drift correction functionality to allow for the calculation of drift based on DAPI images instead of fiducial beads, as well as correcting drift along the z-axis (perpendicular to the microscope slide). Images were aligned across hybridization rounds by identifying local maxima and minima in the DAPI channel and calculating the offset of these anchor points using fast Fourier transform. A high-pass filter and deconvolution using a Gaussian point spread function were applied to reduce noise and background of images, then a vector was constructed for each pixel representing the brightness of that pixel across all imaging rounds. An L2 normalization was applied to these vectors to standardize their total brightness, and then the Euclidean distance to each of the barcodes in the MERFISH codebook was calculated. For this calculation, each barcode was represented as a similar L2-normalized brightness vector with a value of 0 in the off-bit rounds and 1 in the on-bit rounds pre-normalization. Each pixel was assigned the gene identity of the nearest barcode, provided the distance was less than 0.6, which was chosen to capture distances representing single bit errors. Adjacent pixels with the same gene identity were combined as a single RNA molecule. Scale factors to equalize the brightness across imaging rounds were calculated by decoding a random sample of 50 z-slices and keeping RNA molecules with at least 5 pixels, then calculating the factors necessary to equalize the mean brightness of the pixels in these molecules in each round. This process was repeated iteratively 10 times, with brightness in each iteration normalized using the scale factors calculated in the previous iteration. The RNA molecules were filtered based on 3 parameters: the number of pixels in the molecule, the mean brightness of the pixels, and the minimum distance to the barcode among the pixels. A 3-dimensional histogram was constructed using these parameters, and then for each histogram bin, the ratio of gene barcodes to blank barcodes was calculated. The barcodes from the bins with the lowest gene-to-blank were removed until a misidentification rate of 5% was reached.

#### Cell Segmentation

Cell segmentation was performed using the same method as above using Cellpose v2.

#### Cell clustering

The cells were first clustered using the leiden algorithm and manually labeled with major cell types based on gene expression. Clusters with the same major cell type label were merged. Next, each major cell type was sub-clustered individually and manually labeled based on gene expression and the spatial pattern of the cells’ locations. Some clusters were identified as artifacts. Further inspection of these artifact clusters revealed that some were the result of out-of-focus images in specific rounds of imaging, affecting 72,142 (2.3%) cells. Because only a subset of genes were affected by these problematic images, the unaffected genes were used to impute the correct cell type by nearest-neighbor classifier using cells outside the affected regions. Other artifact clusters were identified to consist of low-quality cells arising from segmentation artifacts or low transcript counts and were removed from the dataset.

#### Slice alignment and 3D reconstruction of the human developing heart

In order to reconstruct the 3D map of the human developing heart, we utilized Spateo ^26^ we previously developed and extensively demonstrated its favorable performance when compared to PASTE, PASTE2, Moscot, STAlign, etc.. In brief, Spateo leverages the transcriptional similarity between cells in adjacent slices, together with a spatial constraint on the transformation of the slice, to align the 53 slices of the developing human heart. Specifically, Spateo relies on a Bayesian generative model to efficiently and robustly align the adjacent slices, and to output a probabilistic mapping and a rigid or non-rigid transformation. Although Spateo’s non-rigid alignment mode can eliminate distortions introduced by tissue sectioning, the data has minimal distortion as the cardiac cavities are relatively small and don’t lead to substantial distortions. Additionally, we cannot determine whether the differences between sections are caused by the sectioning process or represent genuine biological variation. Therefore, we opted for Spateo’s rigid alignment mode. Furthermore, due to partial overlap between certain sections (such as sections 21 and 22), we set the partial_robust_level parameter of Spateo’s *morpho_align* function to 25 for these sections to achieve more precise alignment and reconstruction results. For the input gene expression features, we utilized principal component analysis (PCA) features processed by the Scanpy Python package, which provides a denoising effect compared to the original raw counts. Default values are used for all other parameters in Spateo’s *morpho_align* alignment function. The entire reconstruction process iteratively applied Spateo’s alignment method along the z-axis, enabling us to precisely reconstruct a 3D human heart containing 3.1 million cells. All computations were performed on a high-performance server running Ubuntu 22.04.5, equipped with Intel Xeon Platinum 8280 processor, 1.4 TB of RAM, and eight NVIDIA A100 GPUs with 40 GB memory. The complete reconstruction process required approximately 1 hour of processing time.

#### 3D Visualization

After the 3D reconstruction of the human heart, we utilized Spateo’s built-in 3D rendering method for visualization, which is built upon PyVista ^95^ and provides convenient and realistic rendering capabilities. For 3D visualization of cell types, we used Spateo’s *construct_pc* function to convert the h5ad data structure into VTK files. For scenarios displaying 3D scalar values, such as gene expression levels in 3D, we additionally employed Spateo’s *add_model_labels* function to insert the desired scalar values into the VTK files. To address potential occlusion issues, e.g. when some parts of the data (e.g., cells, regions, or structures) block or obscure other parts from view, in 3D visualization, we rescaled the scalar values to be between 0 to 1 range and use them as opacity values. Finally, we used Spateo’s *three_d_plot* function for 3D rendering.

#### Density measurement of each cell type

For different cell types, we calculated their *k*-nearest neighbor density to more accurately capture the local density distribution of cells. Specifically, for each cell, we first calculated the spatial distance *d* to its *k*-th nearest neighbor of the same cell type (where *k=30* in this work), then constructed a sphere with radius *d*. The density was then defined as the number of cells of the same cell type within this sphere divided by the volume of the sphere, where the volume is defined as 4/3 π*r*^3^.

#### Cell-type co-existence neighborhood analysis

This method was used to generate **Figures 5A and 5B**. We aimed to identify pairs of cell types that are more likely to co-exist, thereby potentially uncovering spatial relationships between different cell types. To this end, we first constructed cell neighborhoods according to the definition described in the “Neighborhood complexity calculation” section and calculated the proportional composition of cell types. Given *N* cells and *M* cell types, we obtained an *NxM* matrix, where the *i*-th row represents the proportional composition of cell types in the neighborhood of the *i*-th cell. From a column perspective, we obtained the distribution of the proportional composition of each cell type across all cell neighborhoods, described by an *Nx1* vector. Then, for any two different cell types *m1* and *m2*, we calculated their Pearson correlation coefficient between two *Nx1* vectors, which represents a co-existence score between cell types. A high Pearson correlation coefficient between two cell types indicates that these cell types tend to either co-exist or be absent together in most neighborhoods. To further discover clusters of spatial co-existence patterns of cell types, we constructed a dendrogram structure based on the cell type correlation coefficient matrix. More specifically, we first calculated the pairwise similarity between cell types using Scipy’s ^96^ *pdist* function with the “cityblock” metric. Then, we constructed the dendrogram structure using scipy’s linkage function with the “single” method. To visualize the co-existence relationships between cell types more intuitively, we further utilized the igraph Python library to visualize the network relationships between cell types. In the network, nodes represent different cell types, and edges are undirected with weights representing co-existence scores. To reduce spurious correlations, we filtered out connections with co-existence scores below 0.07. We manually annotate the dendrogram with five major niche neighborhoods.

#### Neighborhood celltype enrichment analysis

This method was used to generate **Figure 5H**. Leveraging our comprehensive 3D spatial information, we conducted an in-depth analysis of cell-type distributions and their spatial relationships within the human heart. We examined the cellular organization along the *Apex-Base* axis. Specifically, we analyzed the cellular composition within a 30 μm radius for three long branches of the coronary arteries (LAD, LCX, and RCA). The enrichment levels were quantified by normalizing against background cell-type frequencies in the whole 3D heart, revealing distinct patterns of spatial organization along this anatomical axis.

#### Constructing artery vessels network

This method was used to generate **Figure 5E**. We initially identified coronary vessel cells based on *MYH11* enrichment. Specifically, the top 3% of cells with the highest MYH11 expression were selected as a rough set of coronary vessel cells. Cells belonging to the aorta (AO) and pulmonary artery (PA) were then excluded according to their spatial distribution. To further refine the selection, we applied spatial density estimation to filter out randomly scattered outliers (performed in three iterations, removing the lowest-density 10%, 3%, and 0.5% of cells, respectively, see section “*Density measurement of each cell type*”). This procedure yielded a high-quality set of coronary vessel cells.

To identify each coronary vessel cross-section, we implemented a spatial clustering method using the Density-Based Spatial Clustering of Applications with Noise ^97^ (DBSCAN) algorithm from the scikit-learn library. This approach enabled us to identify and segment distinct vessel cross-sections within each tissue slice, which were subsequently designated as nodes in our vascular tree structure. DBSCAN parameters were empirically optimized, with the neighborhood radius (*eps*) set to 50 μm and the minimum cluster size (*min_samples*) set to 5. To ensure robust vessel identification, we excluded noise points (cells assigned with label −1) from the analysis.

Leveraging the inherent hierarchical, top-down organization of the cardiac surface vasculature, we developed a top-down tree construction algorithm. For each vessel node identified in slice *z*, we established hierarchical connections by computing a connection loss between the current node and each visited node, defined as:

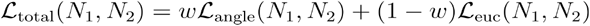

where the angular and Euclidean terms are:

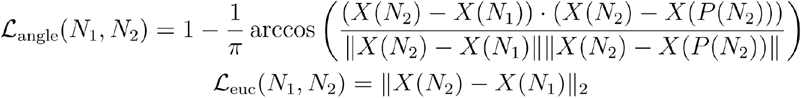

Here *N*_1_ denotes the current node, *N*_2_ a visited node, *X*(·) the centroid of cells in a node, and *P*(·) the parent of a node. The angular loss ensures the connected nodes across three consecutive layers are on a relatively straight line while the second Euclidean loss ensures the distance between two nodes (*N*_1_, *N*_2_) is minimal. The hyperparameter w balances the contributions of the two terms and was set to 0.5. At each step, the current node was assigned to the visited node with the lowest total connection loss, after which the current node was added to the visited set. To maintain biological plausibility, node connections within the same section were disallowed, and only connections to nodes in higher layers were permitted. In cases where the top-down relationship is not strictly satisfied (e.g., at the top of the LCX, where the vessel initially ascends before descending), predefined manual connections were introduced to preserve vascular continuity.

#### Thickness calculation for heart layer

This method was used to generate **Figures 4H-4K** and **S8**. To quantify the 3D thickness between different layers, we first reconstructed the surface geometry of each layer using a mesh-based representation. Since the interventricular septum (IVS) region may affect the thickness estimation and is not of our primary interest, we subsequently identified and removed this region. Finally, based on the reconstructed meshes, we computed the physical 3D thickness between layers. The detailed steps are described as follows.

#### Construction of Layer Surfaces

This method was used to generate **Figure 4H-4K**. We began by defining the cell types contained within each layer to construct a dense, non-hollow point cloud. For instance, the mesh of the epicardial layer was supported by both left and right ventricular cardiomyocytes (vCMs) together with epicardial cells. To obtain a spatially compact and continuous cell distribution, we performed density-based filtering of the outlier cells. Specifically, for cell types with left–right distinctions, we manually separated left and right cells and then applied the density filtering procedure described in Section “Density measurement of each cell type”. We iteratively removed the lowest 1% of low-density cells until the remaining cells occupied a spatially continuous region.

Next, we reconstructed the surface mesh using the *st*.*tdr*.*construct_surface function* in Spateo, employing the marching-cubes algorithm with parameters “*mc_scale_factor*”: 1.05, ‘*dist_sample_num*’: 1000, and ‘*levelset*’: 0.4. Because the spacing between slices was relatively large, we compressed the z-axis of the point cloud to one-fifth of its original scale during mesh construction, and subsequently rescaled it fivefold to restore the true physical z-axis dimension. This approach effectively prevented the generation of internal voids. Using this procedure, we reconstructed three major layers: epicardial, endocardial, and hybrid layers.

#### Identification of the IVS Region

This method was used to generate **Figure 4H**. The IVS region is generally located between the left and right endocardial layers. Thus, we first identified the IVS on the endocardial meshes. For the left endocardial mesh, we computed the normal vector at each vertex and determined whether the normal intersected the right endocardial mesh. If such an intersection existed, the vertex was labeled as belonging to the IVS region. The same procedure was applied symmetrically to the right endocardial mesh.

After identifying IVS regions on the endocardial surfaces, we further detected IVS regions on the epicardial mesh. Again, we used the normal-intersection method and classified the intersection patterns into five cases: (1) The normal vector did not intersect any endocardial mesh. (2) The normal vector intersected the right endocardial mesh, and the intersection point was labeled as part of the IVS. (3) The normal vector intersected the left endocardial mesh, and the intersection point was labeled as part of the IVS. (4) The normal vector intersected the right endocardial mesh but was labeled as a non-IVS region. (5)The normal vector intersected the left endocardial mesh but was labeled as a non-IVS region. We defined cases (1)–(3) as IVS regions, while cases (4) and (5) were assigned to the right and left walls, respectively.

#### Computation of 3D Physical Thickness

This method was used to generate **Figures 4H-4K**. Based on the reconstructed layer meshes, we calculated the 3D physical thickness within the non-IVS regions on the epicardial surface. For each vertex on the epicardial mesh, we first computed its surface normal vector and then determined the intersection points of this normal with the endocardial, hybrid, and Purkinje layer meshes. The distances between consecutive intersection points along the normal direction were defined as local layer thicknesses.

Specifically, the distance between the Purkinje layer and the endocardial layer was defined as the trabecular layer thickness, the distance between the endocardial layer and the hybrid layer was defined as the hybrid layer thickness, and the distance between the hybrid layer and the epicardial layer was defined as the compact layer thickness. The distance between the epicardium and the endocardial layer was defined as the free wall thickness. This approach minimizes the inclusion of free trabecular muscle in the measurement of ventricular wall thickness, as these structures can vary with the contractile state of the heart at the time of dissection. For instance, during contraction, free trabeculae may cluster together and be mistakenly identified as part of the compact ventricular wall. This pointwise computation provided a spatially resolved map of heart layer thickness, enabling quantitative comparison across different regions of the heart.

#### Gene expression gradient analysis

This method was used to generate **Figures 5I-5K**. Based on the hierarchical tree structure of the vasculature, we identified the path from each vascular cell to its corresponding root of the vessel. We can then perform path-dependent differential gene expression analyses to discover gene expression gradient along the vascular hierarchy.

#### Correlation of RNA expression measurements

All pseudo-bulk correlations shown in this work are Pearson correlations calculated using the *log*_*10*_(x + 1) transformed mean transcript count per gene across cells based on the raw cell by gene tables (UMI count for scRNA-seq or localized transcript count for MERFISH and smFISH). Cell type to cell type correlation heatmaps were generated by first calculating the mean transcript count per gene among the cells of each cell type, then transforming each mean into a z-score using the distribution of means for each gene across cell types. The Pearson correlations of these z-scores between every pair of cell types were used for the heatmap values, and the average of the maximum correlation for each cell type to any other cell type was used to quantify the overall correlation.

### Spateo-VI integrates various 2D and 3D spatial transcriptomics and other modalities

#### Spatial Graph Construction

This method was used to generate **Figure 6**. The illustration of the method is shown in **Figure 6A**. To leverage spatial context, we construct a graph of cell neighborhoods based on each cell’s spatial coordinates. We provide two options for graph construction: K-nearest neighbors (KNN) and Delaunay triangulation. In KNN mode, each cell is connected to its *k* nearest neighboring cells in space, yielding an adjacency graph that captures local spatial proximity. In Delaunay mode, we perform Delaunay triangulation on the coordinates, connecting cells that share a triangle edge, which tends to capture both nearest and slightly farther neighbors in a planar arrangement. These options are configurable (e.g. *spatial_graph_type=“knn” or “delaunay”*). For KNN, we use a default of 10 neighbors (or fewer if fewer cells) and ensure the graph is symmetric (if cell *i* is among cell *j*’s neighbors, we add the reciprocal edge). For Delaunay, we triangulate the points (adding a small jitter if points are collinear or 1D) and connect all cells that form the vertices of a common Delaunay simplex. The result in both cases is an undirected graph *G* = (*V, E*) with nodes as cells and edges representing spatial adjacency. If the graph happens to be disjoint or have isolated nodes, we fall back to a fully-connected or KNN graph to ensure all cells are reachable. The spatial graph construction is a preprocessing step that encodes spatial structure and serves as a prior for information sharing between neighboring cells for downstream use in the model’s inference process.

#### Spatial Graph Attention-based Encoder

Given the spatial neighbor graph, our model employs a Graph Attention Network (GAT) to incorporate spatial information into the cell representations. Starting from each cell’s initial latent representation (described below), we apply a graph attention layer that propagates information across the neighbors of each cell. Specifically, we use a multi-head GATv2 convolution layer, which allows each cell to attend to its neighbors with learned attention weights. Formally, for a cell *i* with latent vector *z*_*i*_, and neighbors *j* ϵ *N*(*i*)on the spatial graph, the graph attention layer computes a new feature *h*_*i*_ as a function of the weighted aggregate of neighbor features:

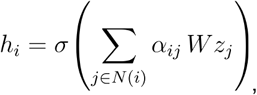

where *W* is a learnable weight matrix and *α*_*ij*_ is the attention coefficient for neighbor *j* of *i*. The attention coefficients *α*_*ij*_ are computed via a softmax that considers *i*’s neighbors’ features, enabling the network to assign different importance to different neighbors. Multi-head attention to multiple aspects of the neighborhood, and the outputs from multiple heads are averaged. This GAT-based message passing effectively smooths and enriches each cell’s latent representation with information from its spatial vicinity, while automatically learning the relevance of each neighbor.

The output of the graph attention layer for cell *i* is a spatial feature vector *h*_*i*_ of dimension *n*_spatial_. We interpret this as parameters of a latent spatial factor for each cell. In particular, we assume that for each cell *i* there is an unobserved spatial latent vector 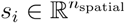 that follows a Gaussian distribution conditioned on *h*_*i*_. We use two parallel linear layers to produce a mean 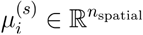 and log-variance 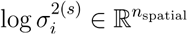 from the GAT output[4]. This yields a variational posterior for the spatial latent:

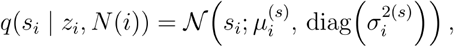

where 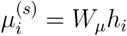, and 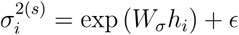, and (with a *ϵ*=10^−4^small for numerical stability). In practice, is sampled by the reparameterization trick as 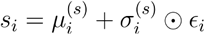 with 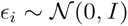. Intuitively, this *spatial encoder* looks at a cell’s expression-derived latent vector and its neighbors’ latents, and then proposes a distribution for a spatial effect *s*_*i*_ for that cell. If no neighbors are present or the graph is empty, the encoder simply yields a default *s*_*i*_ ≈ 0 with unit variance, meaning no spatial adjustment. By using attention, the model can learn to ignore irrelevant neighbors or focus on certain directions (e.g. along a tissue boundary) as needed.

#### Variational Inference and Objective

Our model follows the single-cell variational inference (scVI)^74^ framework, using a variational autoencoder (VAE)^98^ to model gene expression data. Each cell *i* has an underlying global latent variable 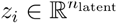 that captures its transcriptional state (accounting for technical effects like batch, if specified). We assume a standard normal prior 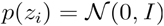. An encoder neural network (a multilayer perceptron with *n*_layers_ hidden layers and *n*_hidden_ nodes each) takes the observed gene expression vector (and optionally other covariates) and outputs the approximate posterior distribution 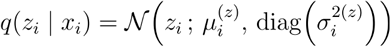. This is analogous to standard scVI: the encoder learns to compress the high-dimensional gene expression of a cell into a lower-dimensional latent representation. In our Spateo-VI model, if spatial information is used, the inference proceeds in two stages: first infer *z*_*i*_ from the cell’s own data, then infer *s*_*i*_ from *z*_*i*_ and neighbors as described above. Thus, the overall approximate posterior factorizes as 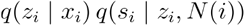. We emphasize that *s*_*i*_ is only inferred if use_spatial=True and it encodes residual variation structured by spatial locality.

On the generative side, the model reconstructs the observed gene expression from the latent variables. We model raw gene counts (such as UMI counts) with appropriate count distributions commonly used in single-cell analysis. By default we use a negative binomial (NB) likelihood (or zero-inflated NB if specified) for each gene’s expression, which accounts for over-dispersion and the many zero counts. Specifically, for each gene *j* in cell *i*, our decoder outputs a parameter *μ*_*ij*_ (the mean of the NB) and gene-specific overdispersion *θ*_*j*_. The mean is formulated to capture library size and latent effects: *μ*_*ij*_ = *𝓁*_*i*_ · *f*_*j*_(*z*_*i*,,_ *s*_*i*_), where *l*_*i*_ is the cell’s total count (library size) or an estimated size factor, and *f*_*j*_(*z*_*i*,,_ *s*_*i*_) is the decoder function’s output for gene *j*. In practice *f*(·) is a neural decoder that takes *z*_*i*_ (and potentially *s*_*i*_ or other covariates) and produces gene-wise rates; when no spatial latent is used, *f* only depends on *z*_*i*_.

Each gene count is then modeled as 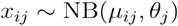. The negative log-likelihood (reconstruction loss) for gene *j* in cell *i* is:

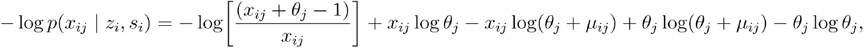

which is the standard NB formulation (for zero-inflated NB, an additional zero-inflation probability is learned per gene). The decoder is trained to maximize the likelihood of the observed counts given the latent variables.

The overall training objective is the evidence lower bound (ELBO) from variational inference. For a single cell *i*, the ELBO is:

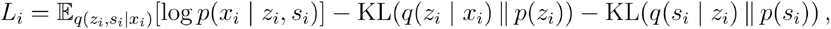

where *p*(*s*_*i*_) = *𝒩* (0,*I*)is the prior for the spatial latent. In practice, we introduce a weighting factor ω for the spatial KL term to moderate its influence: the objective becomes:

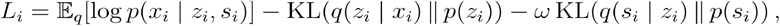

The hyperparameter ω corresponds to *spatial_kl_weight* in our implementation (default ω = 0.01). The KL divergences have closed forms since *q* and *p* are Gaussians. During training, we average *L*_*i*_ over all cells and maximize it. We use stochastic optimization with Adam, as in standard scVI. Gradients flow through both the encoder for and the GAT-based encoder for *s*. The inclusion of the spatial graph and latent does not drastically increase computational complexity; the KNN graph can be precomputed efficiently, and the GAT convolution is applied on latent dimensions. Graph operations are accelerated via the PyTorch Geometric package.

After training, the model can output various representations: the global latent embeddings *Z* ={*z*_*i*_} for all cells (useful for visualization or clustering), as well as spatial latent embeddings *S* ={*s*_*i*_} which capture spatially correlated variation. We can also impute or denoise gene expression, generate samples, or perform differential expression by leveraging the learned generative model, similar to scVI’s downstream applications.

#### Multi-Modal Data Integration

A key feature of our Spateo-VI model is the ability to jointly model multiple data modalities in a shared latent space. In many experiments, one might have spatial transcriptomics data (e.g. MERFISH with spatial coordinates) as well as protein expression data or epigenetics histone modification. Our model extends the single-modality VAE to handle spatial gene expression matrix and an optional protein expression/Histone modification matrix. We maintain a single latent variable per cell that captures the cell’s state in a way that is common to all modalities. Cells from different modalities are all embedded in this same latent space, allowing direct integration and comparison. We instantiate separate encoders for each modality’s observed data, but they map to the same type of latent *z* distribution. Likewise, for decoding, we have modality-specific decoders: one decodes *z*_*i*_ (and *s*_*i*_ for spatial cells) to gene counts in the spatial transcriptomics, another decodes histone modification. In such cases, the decoder for proteins incorporates a background noise model per protein/histone, similar to how TotalVI^99^ models protein counts as a mixture of a specific foreground signal and an unspecific background component. These parameters would be learned by maximizing the protein likelihood.

Overall, by integrating multiple modalities, Spateo-VI can learn a unified latent representation that synthesizes information from spatial gene expression and histone markers. This shared latent space facilitates dataset integration, as well as multimodal downstream analyses. For example, one can cluster cells based on the latent space that reflects both RNA and histone. Our approach is analogous to TotalVI’s joint modeling of RNA and proteins, but extended to spatial transcriptomics and epigenetics: we represent all modalities probabilistically and use a variational inference scheme to decompose observed variation into a common biological latent variable, modality-specific technical effects (e.g. histone background, batch effects), and in the case of spatial data, a spatial effect *s*. This provides a cohesive solution for multi-modal single-cell data integration, allowing tasks such as dimensionality reduction, data integration across modalities, and differential expression to be performed in a unified framework.

#### Spatial cell-cell interaction analysis

This method was used to generate **Figure 6I and S9C**. Furthermore, to investigate spatial cell interactions, we extended COMMOT^75^ by developing the COMMOT-3D algorithm, enabling its application in 3D spaces and accelerating the optimal transport calculation via GPU acceleration. We utilized the CellChat^100^ database to compute the spatial communication strength of each signal on every cell.

To analyze cell–cell interactions on the vascular structures, we focused on three major vessels, i.e., LAD, RCA, and LCX, together with their 30-µm neighborhoods, and performed interaction inference using COMMOT-3D. Subsequently, for each vessel, we selected morphologically intact vessel sections that form complete ring-like vascular structures from top, middle, and bottom segments, and integrated them into a unified spatial coordinates. Specifically, we applied the non-rigid alignment algorithm in Spateo for the alignment and integration. Finally, we visualized the resulting cell–cell interaction flow vector field on the aligned vascular structures.

#### MERFISHEYES Visualization Implementation

The MERFISHEYES website delivers fast and reliable visualization of millions of cells within a web browser. The rendering is optimized through WebGL (Khronos Group. (2009–2025). *WebGL* (https://www.khronos.org/webgl/) in combination with the Three.js framework (Dirksen, R., Cabello, R., & Contributors. (2010–2025). *Three*.*js* (https://threejs.org/) enabled responsive hovering interactions, fast performance, and utilization across desktop and mobile devices. The backend is implemented using the Rust Rocket (Rust Project Developers. (2010–2025). *Rocket: A web framework for Rust* (https://rocket.rs/) web api is used with file compression to reduce latency and server load.

### Chromatin tracing analysis

a. Localization of Fluorescent Spots To calculate fluorescent spot localizations for chromatin tracing data, we followed the following computational steps:
  1. We computed a point spread function (PSF) for our microscope and a median image across all fields of view for each color channel based on the first round of imaging to be used for homogenizing the illumination across the field of view (called flat-field correction).
  2. To identify fluorescent spots, the images were flat-field corrected, deconvoluted with the custom PSF, and then local maxima were computed on the resulting images. A flat-field correction was done for each color channel separately.
b. Image registration and selection of chromatin traces Imaging registration was performed by aligning the DAPI channel of each image from the same field of view across imaging rounds. First, the local maxima and local minima of the flatfield corrected and deconvolved DAPI signal were calculated. Next, a rigid translation was calculated using a fast Fourier transform to best align the local maxima/minima between imaging rounds. FXYD Coordinates: chr19:35003250-35533250(hg38); FXYD Coordinates: chr19:37547865-38078198 (hs1) Nuclear segmentation was performed on the DAPI signal of the first round of imaging using the Cellpose ^94^ algorithm using the “nuclei” neural network model. Following image registration, chromatin traces were computed from the drift-corrected local maxima of each imaged locus as previously described ^40^.

### Protein Density Quantification

The antibody images were flat field corrected, deconvolved and then registered to the chromatin traces using the DAPI signal as described before. For each chromatin trace, the fluorescent signal of each antibody was sampled at each genomic locus 3D location in each cell.

### Code availability

Custom code used for analyzing MERFISH+ and chromatin tracing datasets in this study are available here:https://github.com/epigen-UCSD/MERFISH_Plus_Paper https://github.com/cfg00/MERFISH_Chromatin_Tracing_2024

Custom code used for 3D reconstruction, visualization, analysis, and imputation in this study are available here: https://github.com/aristoteleo/Human_Heart_MERFISH_Analysis_2024

## Supplemental Figures

**Figure S1.**
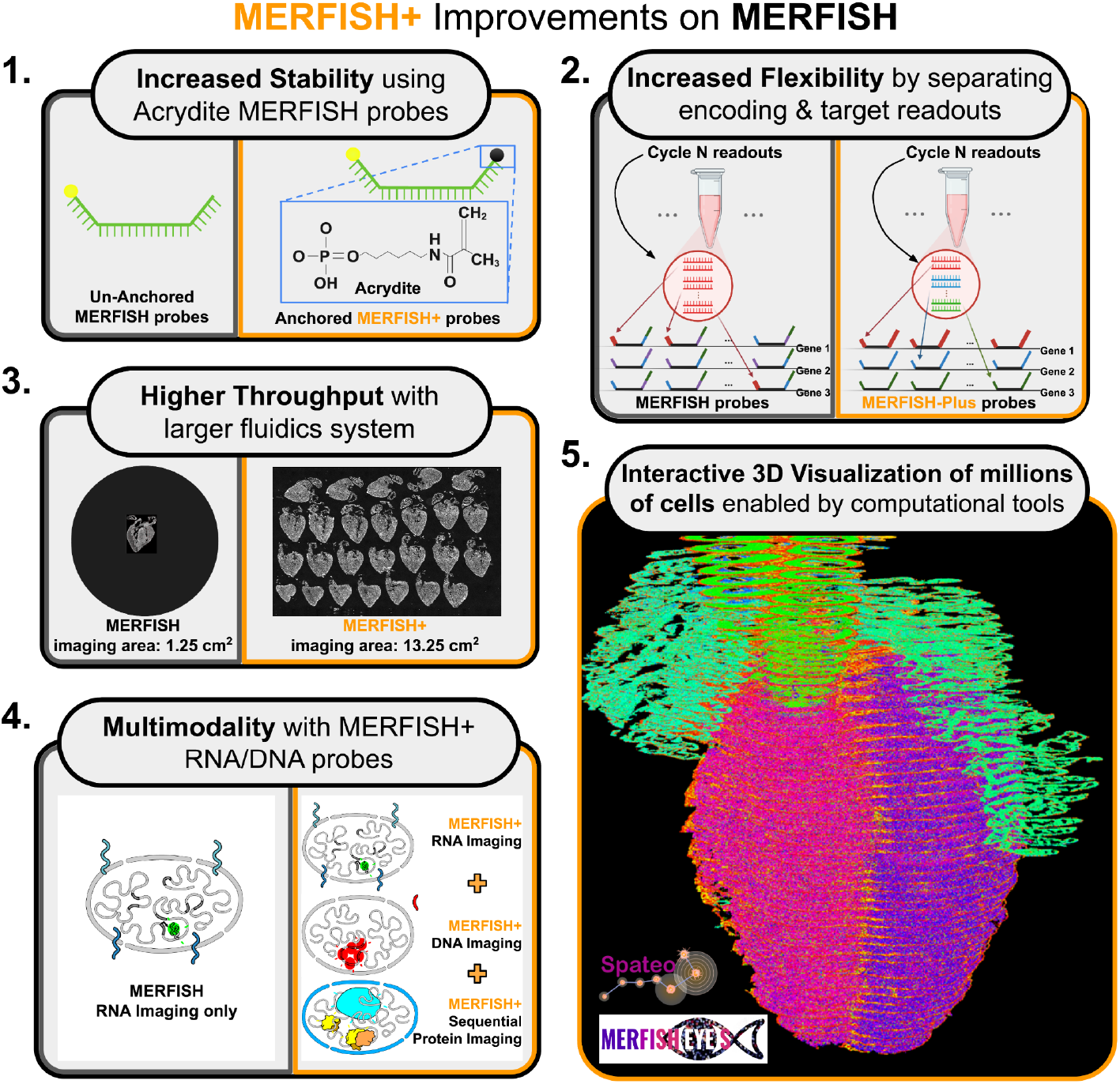
Schematic of the new MERFISH+ capabilities. 1. MERFISH+ probes are synthesized with a 5’ acrydite modification to allow for their robust integration into protective hydrogels. This enhances the stability of experiments. 2. New barcoding strategy of MERFISH+ probes allows for a flexible scaling up of the number of genes profiled. 3. Higher throughput microscopy and microfluidics instrumentation allows for 10X higher imaging area with over 1.5 million cells profiled per experiment. 3. MERFISH+ probes facilitate multimodal and multiplexed imaging of RNA, DNA and histone marks while preserving the integrity of the signal for an extended time. 5. The higher throughput instrumentation, coupled with molecular alignment tools (Spateo-VI) allows for 3D reconstruction of entire human organs.

**Figure S2.**
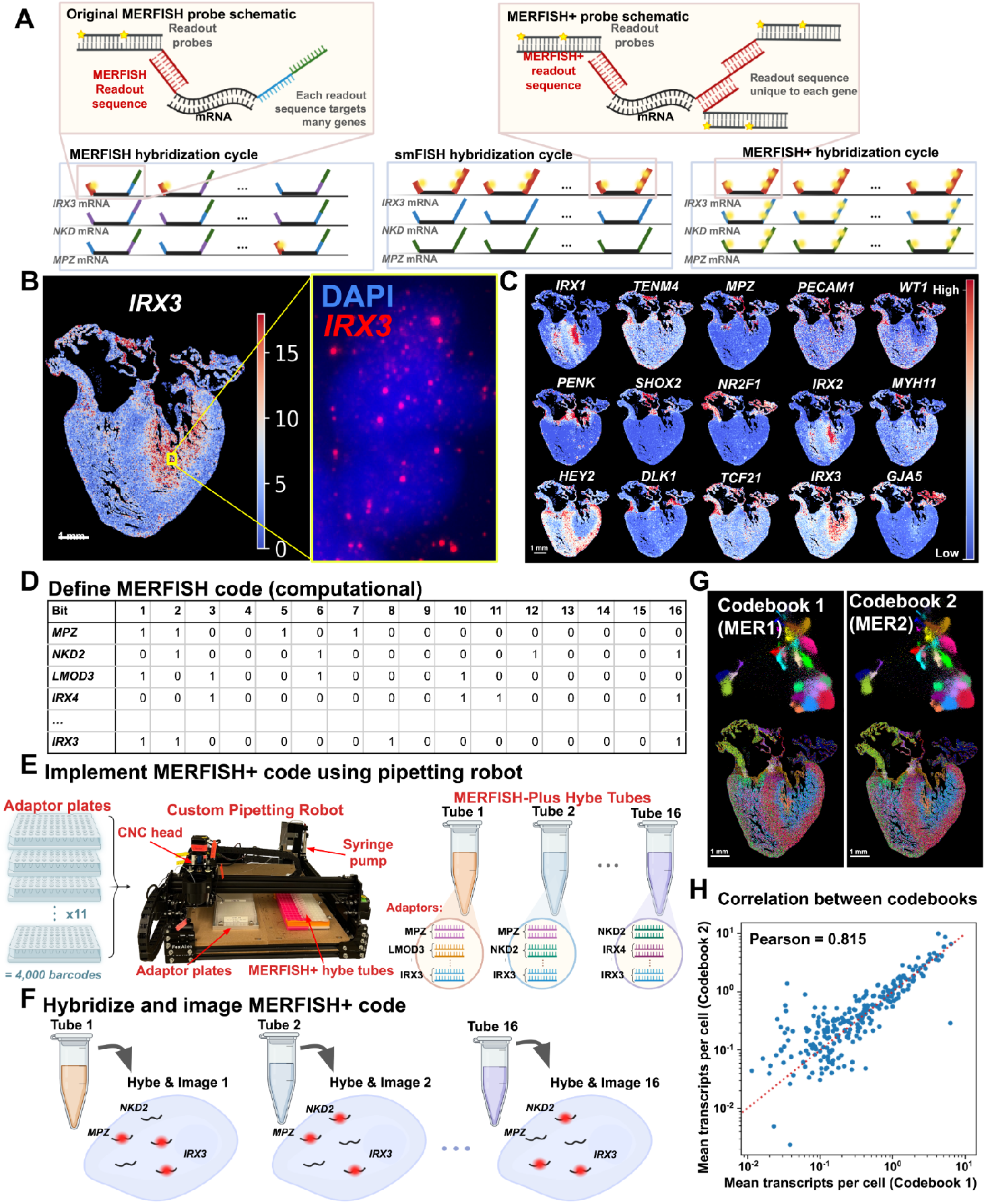
MERFISH+ probes allow for a flexible scaling up of the number of genes profiled. **(A)** The stability of the acrydite-conjugated probes, combined with stringent washing conditions, allow for a redesign of MERFISH probes (termed MERFISH+ probes) in which the readout strategy of the original MERFISH design was changed for increased flexibility. The original MERFISH probe design (**A, top-left**) has built-in readout sequences targeting many different genes simultaneously in each hybridization cycle (**A, bottom-left)**. This design does not enable individual genes to be imaged separately and if any mistakes were introduced during the design (i.e. the accidental inclusion of a gene with high expression), the MERFISH probes need to be reordered and resynthesized. In contrast, the MERFISH+ probe design added built-in unique readout sequences for all probes targeting each gene (**A, top-right**). This allows MERFISH+ probes for any gene or subset of genes to be readout using either through serial single-molecule (sm)-FISH or using MERFISH with an adaptive combinatorial readout strategy (**A, bottom-middle and bottom-right**). **(B) (C)** Single-molecule FISH images of selected genes from a 1,835 MERFISH library hybridized to a 16-um human developing heart section. **(D) (E) (F)** Schematic demonstrating that MERFISH+ probes facilitate designing and imaging custom combinatorial readout strategies for faster quantification of the genes of interest in each cell. An example MERFISH codebook designed for a set of ~250 genes optimized for the human heart is shown in (D). A custom pipetting robot (E) is then programmed to mix combinations of readout probes for each cycle of hybridization. The pipetting program corresponds to a predefined MERFISH combinatorial codebook such that, for instance, a value of 1 in columns 1,2,8 and 16 for gene *IRX3* (D) means that the readout probes for *IRX3* were pipetted and mixed for hybridization cycles 1,2,8 and 16 (F). **(G)** UMAPs and spatial distributions of cell types derived from two MERFISH experiments performed on the same sample and targeting the same genes with two different combinatorial codebooks. **(H)** Correlation of mean transcripts per cell between the two MERFISH experiments in (G) (Pearson’s correlation coefficient of 0.815).

**Figure S3.**
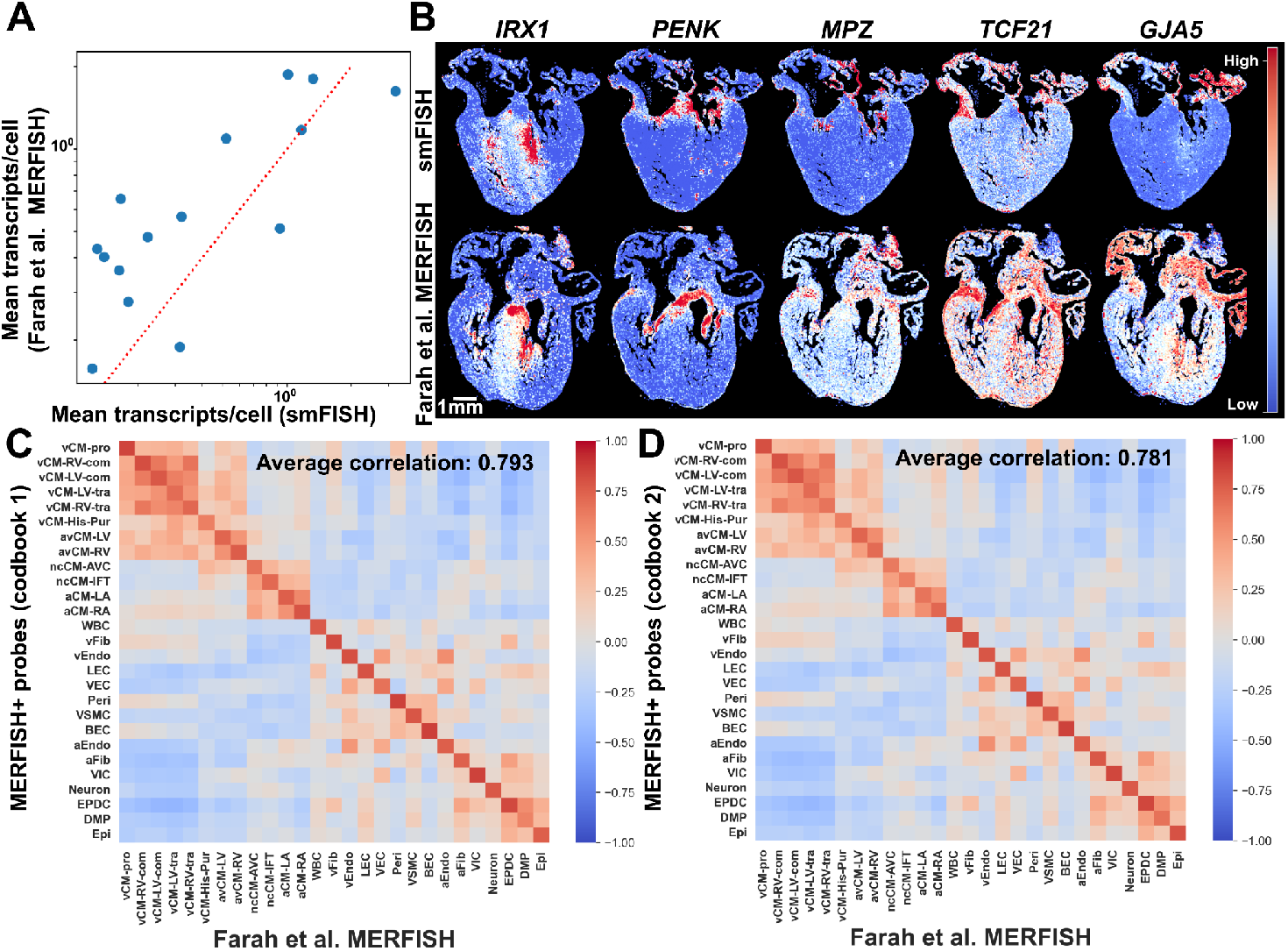
Assessing of MERFISH+ probe performance with prior MERFISH data. **(A)** Correlation of mean transcripts per cell between published MERFISH data^15^ and single-molecule FISH data imaged with the new MERFISH+ probes (Pearson’s correlation coefficient of 0.789). **(B)** Spatial distributions of gene expression per cell comparing published MERFISH data with single-molecule measurements of MERFISH+ probes. **(C, D)** Correlation matrices of average gene expression for 238 genes between MERFISH+ cell-type clusters and prior MERFISH clusters^15^ across 2 codebooks in (C) and (D).

**Figure S4.**
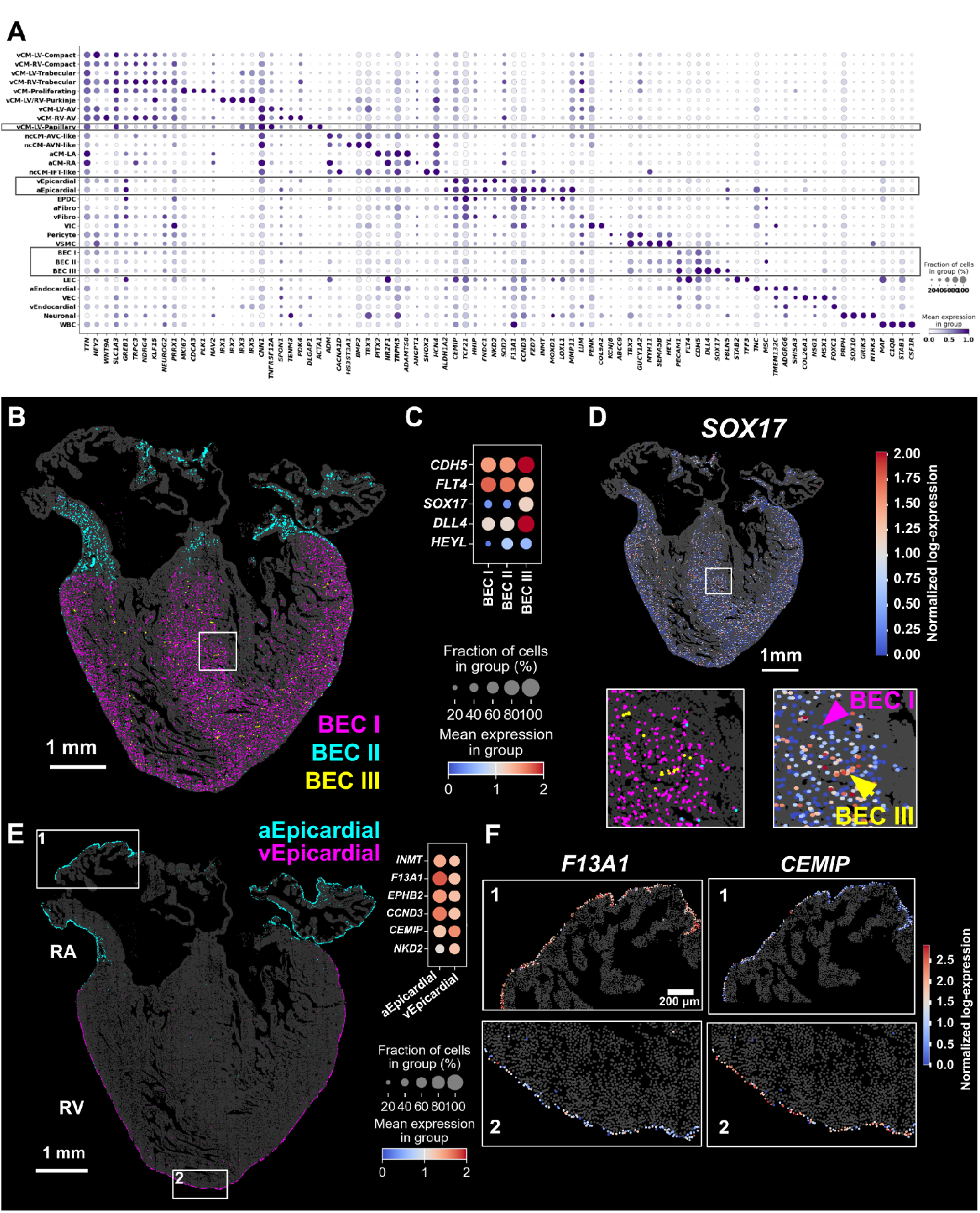
Spatial distribution and gene expression of new cell populations identified with 1,838 genes profiled. **(A)** Dot plot marking the differential gene expression across the cell types defined based on the 1,838 single-cell expression profiles. Newly identified cell populations are boxed. **(B)** Spatial distribution of three BEC sub-populations (BEC I,II and III). **(C)** Gene expression of selected differentially expressed genes across BEC sub-populations. **(D)** Spatial gene expression of *SOX17*, enriched in BEC III cells. Insets show cell type definition of BECs and gene expression of *SOX17* for a zoomed in region of the heart marked by a white box. **(E)** Left: Spatial distribution of aEpicardial and vEpicardial cell populations. Right: Differential gene expression plot across aEpicardial and vEpicardial. **(F)** Spatial gene expression of *F13A1*, enriched in aEpicardial, and *CEMIP*, enriched in vEpicardial.

**Figure S5.**
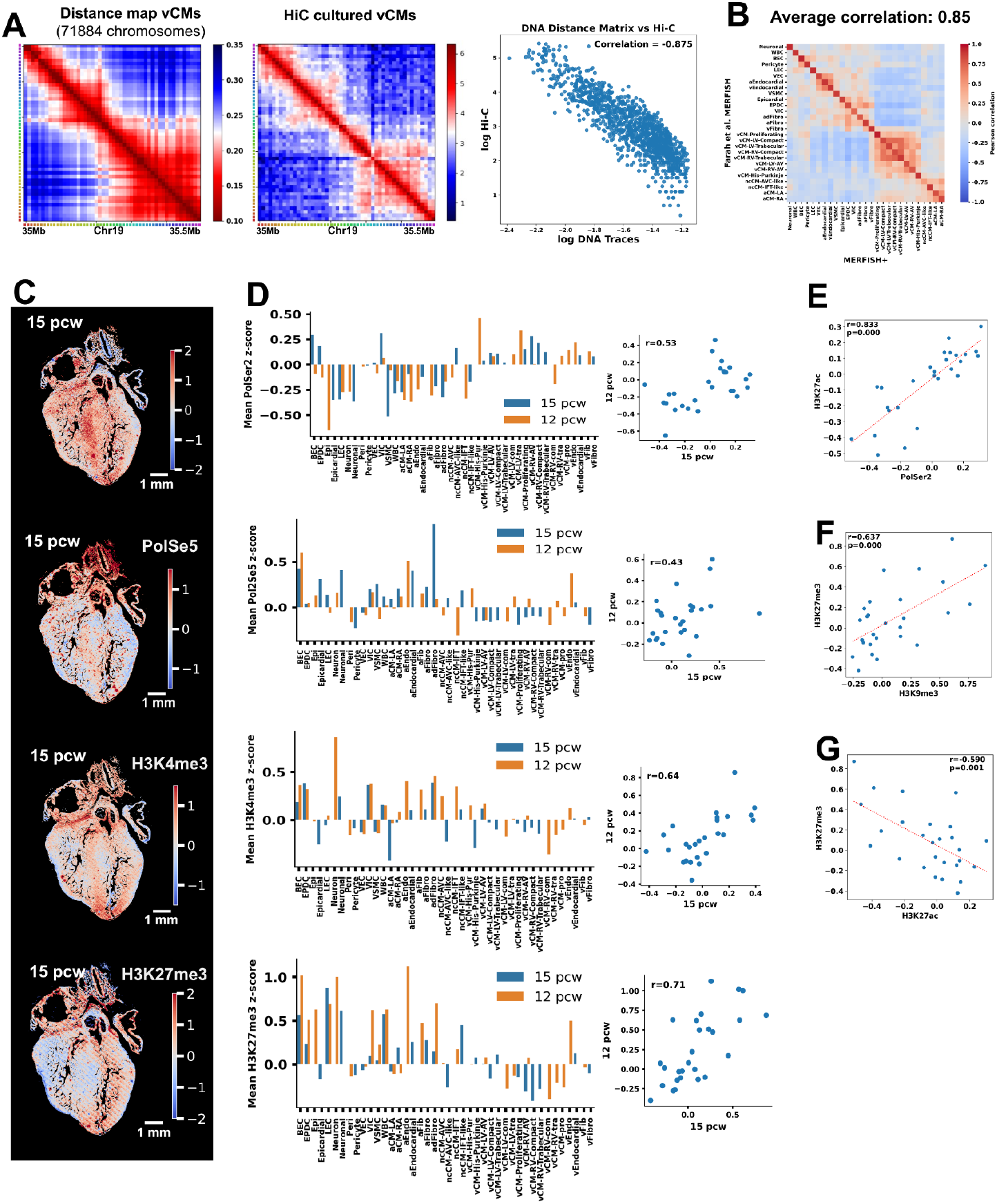
Validation of multimodal imaging results. **(A)** Left: Median distance map across vCMs of the 10kb segments comprising the FXYD locus (chr19:35Mb-35.5Mb) Middle: HiC data^41^ for the FXYD locus in cultured differentiated cardiomyocytes. Right: Correlation of physical distance measured by chromatin tracing in vCMS and the contact probability measured by HiC (Pearson’s correlation coefficient of 0.875). **(B)** Correlation matrices of average gene expression for 238 genes in the multimodal MERFISH+ experiment cell-type clusters and prior MERFISH clusters^15^. **(C)** Spatial map of the z-scored antibody brightness for Pol2ser2, Pol2ser5, H3K4me3, H3K9me3 and H3K27me3 marks across cells in a 15 pcw human heart section. **(D)** Left: Bar plots of z-scored antibody brightnesses of the epigenetic marks in (C) across the cell types of two human heart sections (15 pcw - blue and 12 pcw -orange). Right: Correlation plots of the z-scored antibody brightnesses in (C) across the cell types of the two sections. **(E),(F),(G)** Correlation across cell types in a 15 pcw human heart section for z-scored brightnesses of H3K27ac vs Pol2ser2 (E), H3K27me3 vs H3K9me3 (F) and H3K27ac vs H3K27me3 (G). Pearson correlation coefficients are indicated.

**Figure S6.**
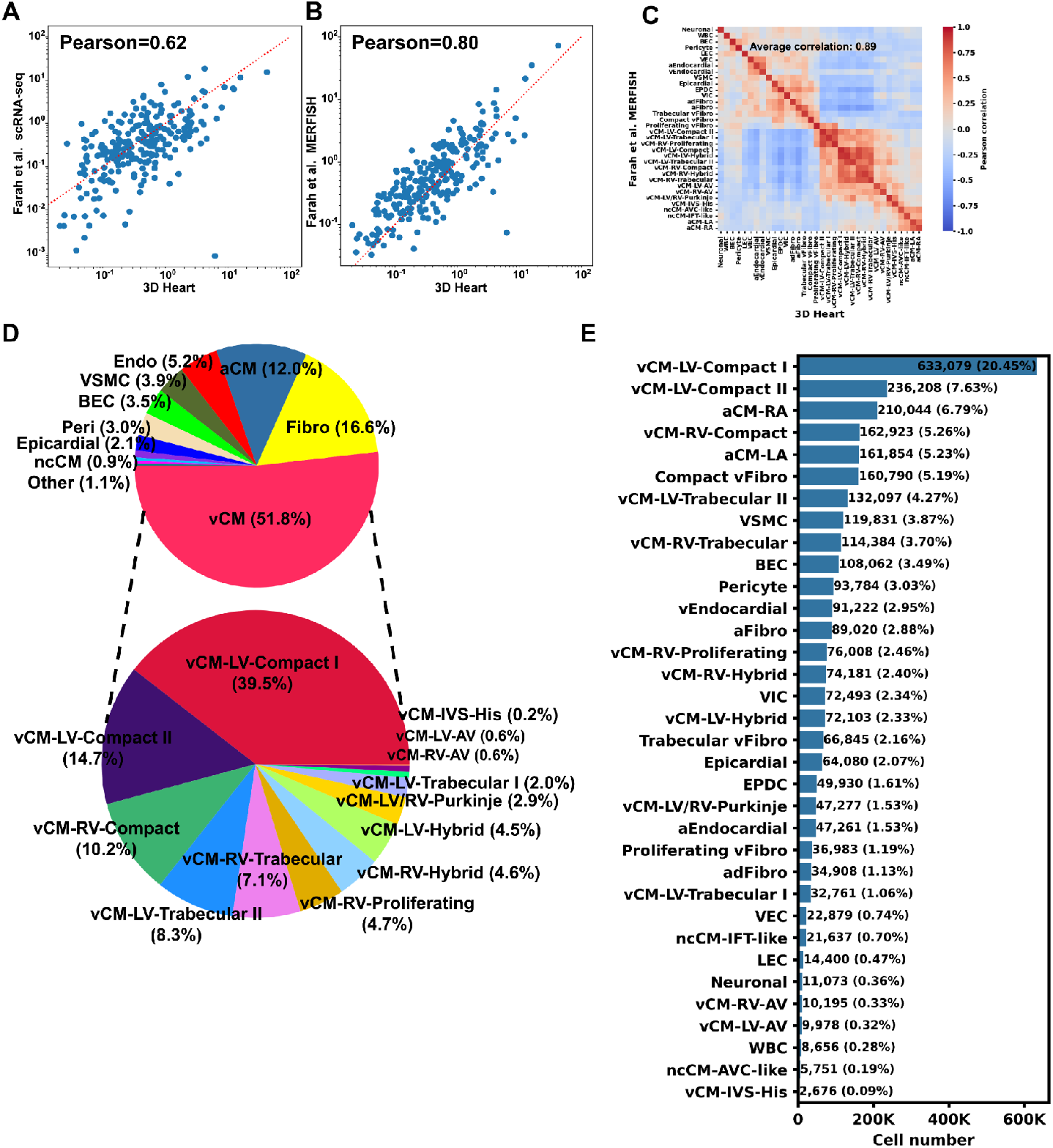
Cell type abundance across the 3D developing human heart. **(A),(B)** Correlation of average transcripts per cell for each gene of the MERFISH+ 3D data with previously published snRNA-seq (A) and previously published MERFISH data (B). **(C)** Heatmap showing pairwise cell type to cell type correlation between this MERFISH data and the previously published MERFISH data. **(D)** Pie chart showing the proportions of major cell types in the heart (top) and the proportions of ventricular cardiomyocyte sub-types (bottom). **(E)** Bar plot with the percentage and number of cells of each cell type across the 3D heart.

**Figure S7.**
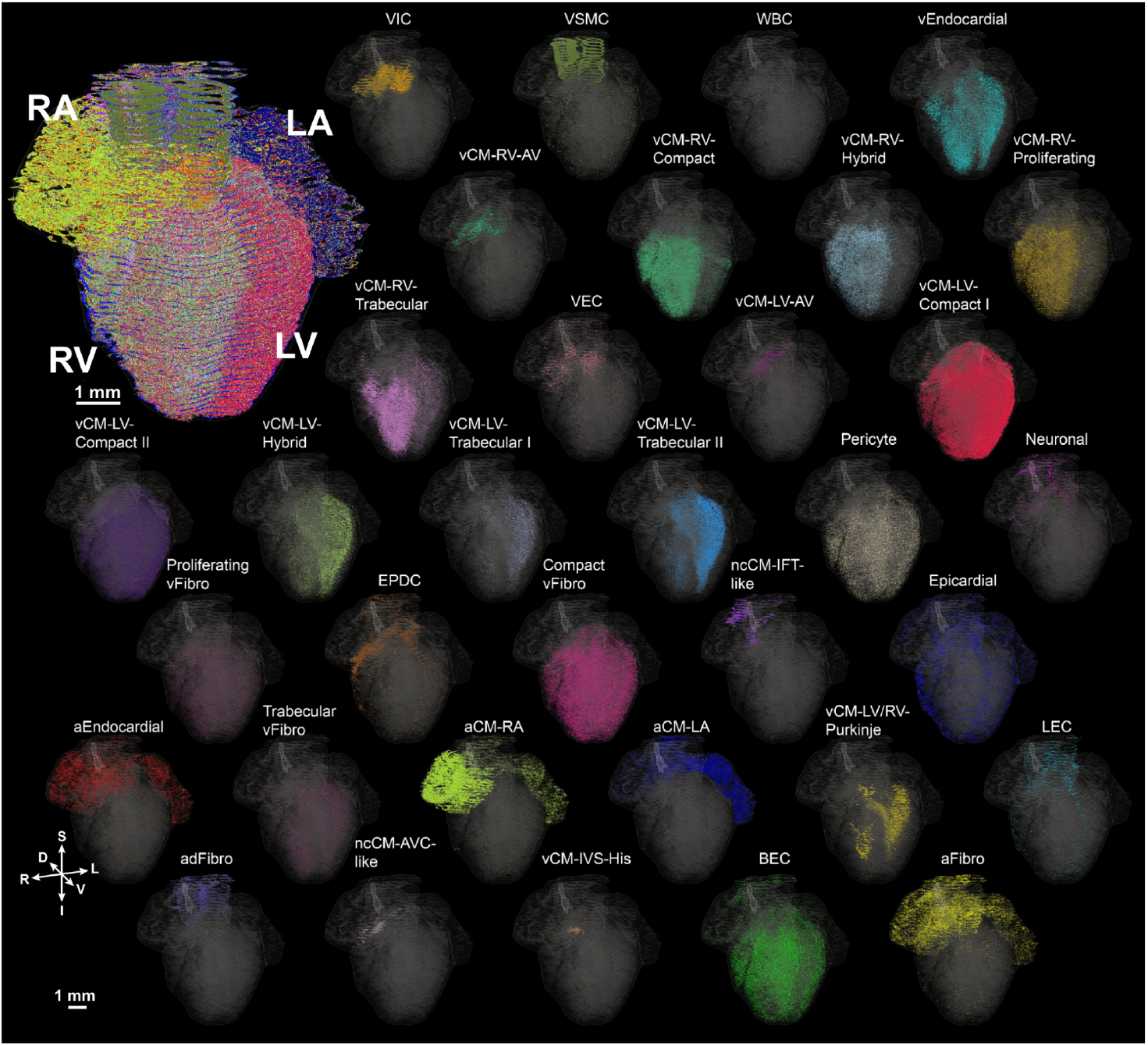
3D spatial distribution of the cell types identified in the developing human heart. A composite image capturing all cell types is shown in the upper left corner. Each subpanel shows the 3D spatial distribution of one of the 34 cell populations identified.

**Figure S8.**
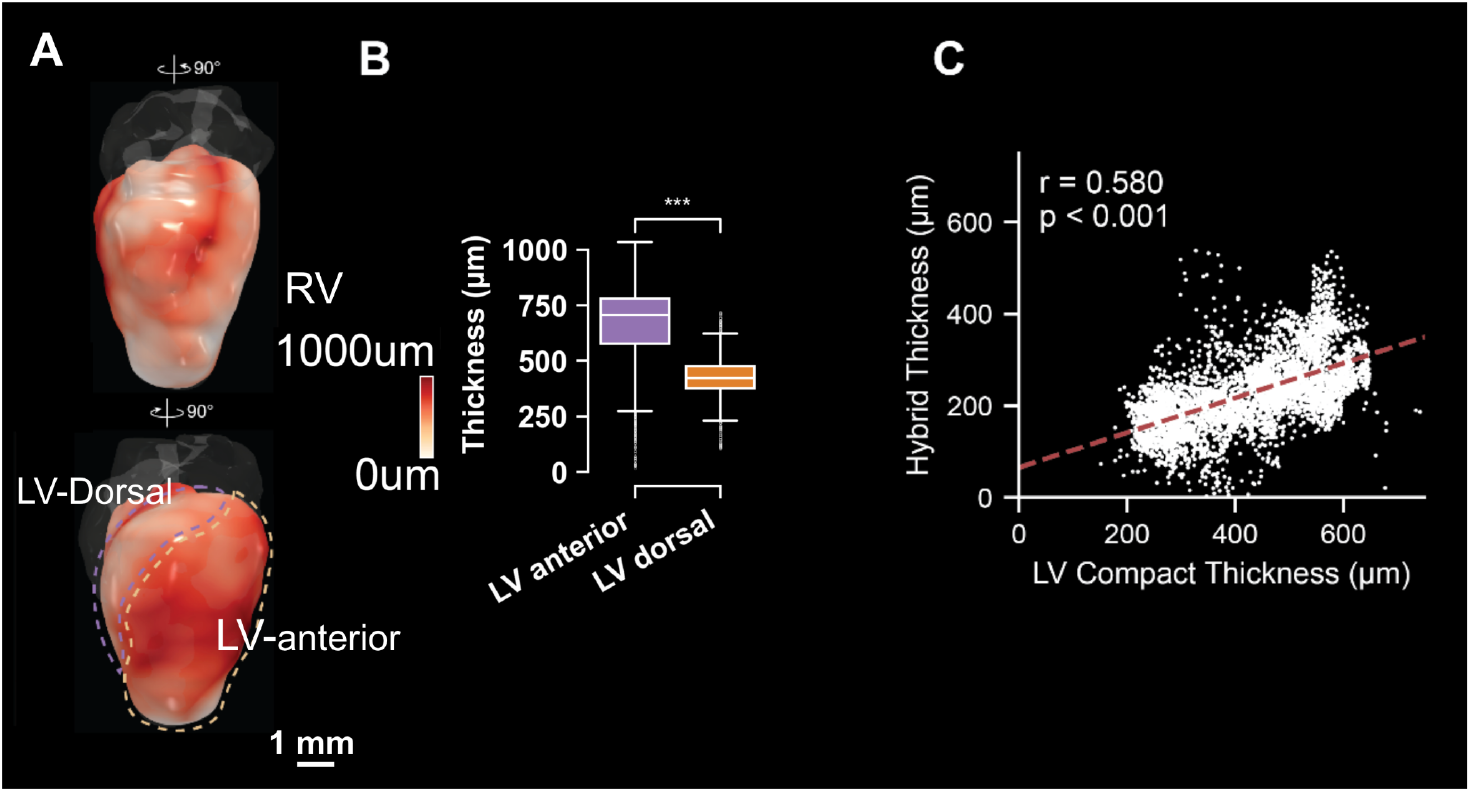
Quantification of the thickness of the ventricular free wall and the ventricular cardiomyocyte layers. **(A)** Thickness of the ventricular free wall measured from the reconstructed 3D heart, displayed across right and left views. The LV-anterior and dorsal, highlighted with dotted circles, where red surface indicates thicker areas and white indicates thinner areas. **(B)** Quantitative comparison of thickness in the LV-anterior and dorsal. *** indicates a p-value < 1e–3 using Student’s t test. **(C)** Correlation of LV hybrid layer thickness and LV compact layer thickness. Pearson correlation coefficient (r) and p-value are indicated. Scale bars as indicated.

**Figure S9.**
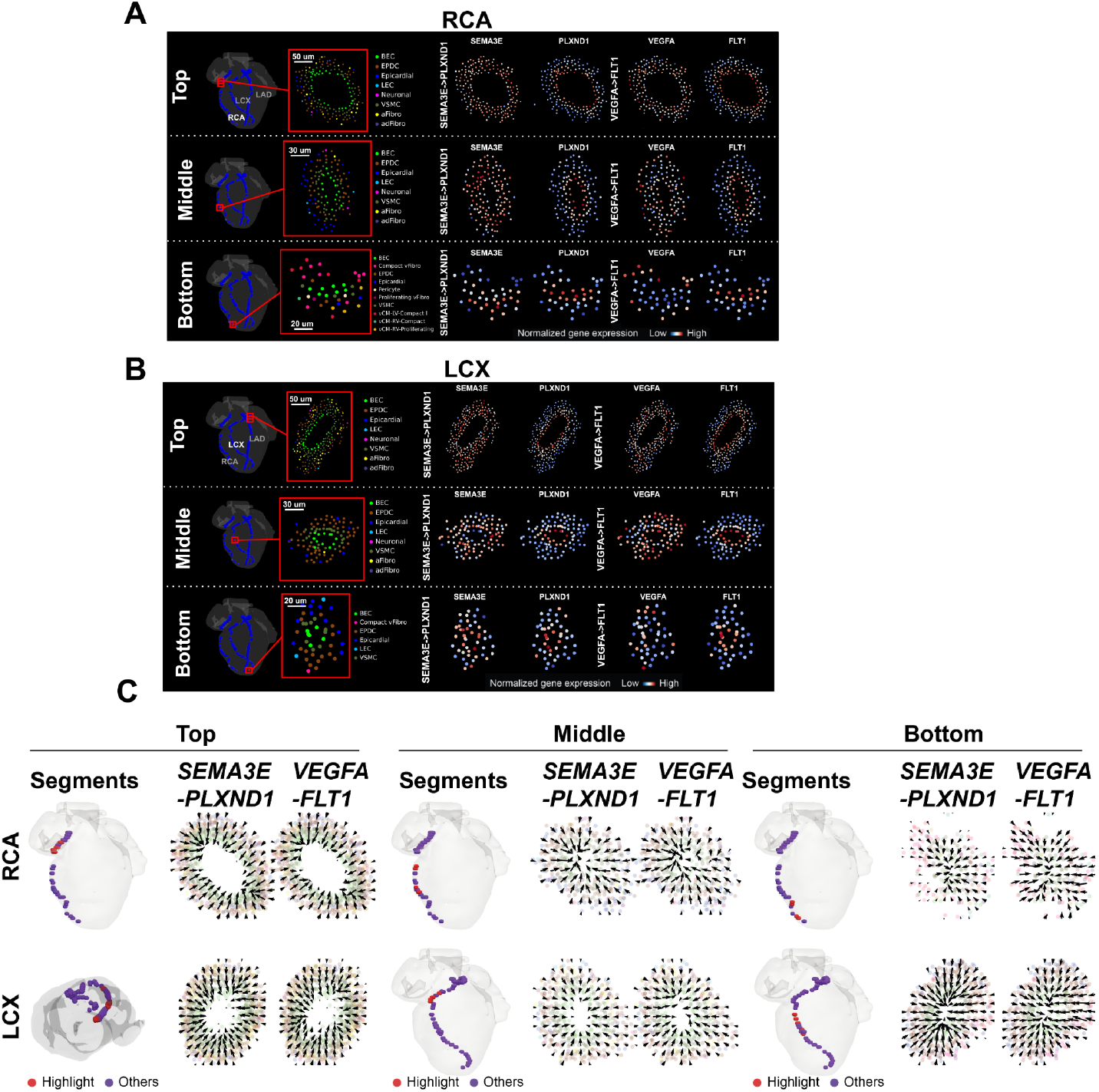
Semaphorin and VEGF signaling in the descending arteries. (**A**),**(B)** Cross-sections of the right coronary artery (RCA) (A) and the left circumflex artery (LCX) (B) showing cell types and imputed gene expression of two known ligand-receptor pairs *SEMA3E*-*PLXND1* and *VEGFA*-*FLT1*. (**C**) Spatial gradient of ligand-receptor interaction in cross-sections of the RCA (top) and LCX (bottom). Three segments, marked in red, along the RCA and LAD from the root to the distal tip are projected to create an aggregate representation of the cross-section. Vector fields (represented by quivers) show the direction and magnitude of *SEMA3E*-*PLXND1* and *VEGFA*–*FLT1* signaling flow in the LCX and RCA.

